## Supplementary material for "The impact of COVID-19 vaccination campaigns accounting for antibody-dependent enhancement": Table S1

**S1 Table.** (Sub-) population sizes of Germany (GER) and the USA chosen in simulations.

| Parameter | Description | GER | USA |
|---|---|---|---|
| $N$ | Total population size | 83 000 000 | 331 000 000 |
| $N^{(\mathrm{NV})}$ | Size of unvaccinable population (40% of $N$) | 33 200 000 | 132 400 000 |
| $N^{(\mathrm{U})}$ | No. of individuals waiting to be vaccinated ($N - N^{(\mathrm{NV})}$) | 49 800 000 | 198 600 000 |
