## Supplementary material for "The impact of COVID-19 vaccination campaigns accounting for antibody-dependent enhancement": S2 Simulations for the USA

### Results for the USA

To illustrate the model's applicability to other countries, we fitted it to the USA as an additional example. The results are qualitatively similar, but differ quantitatively. We will describe the results for the USA in comparison to Germany.

The model parameters are listed in S1 Table - S5 Table and S7 Table.

The average durations for the disease phases ($D_E$, $D_P$, $D_I$, $D_L$ days) and the relative contagiousness ($c_P = c_L$) were chosen identical to those in Germany, as these are parameters describing properties of the virus. The same applies to the Individuals that percentage $f_{\mathrm{Sick}} = 58\%$ of infections that became symptomatic if immunity was not mediated by the vaccine (see S4 Table). Individuals with partial immunity or ADE were again assumed to be half as susceptible to SARS-CoV-2 as unimmunized individuals ($p_{\mathrm{PI}} = p_{\mathrm{ADE}} = 0.5$).

The population size in the USA was set to $N = 331$ million individuals. The first cases were assumed to occur earlier than in Germany, with day $t = 0$ corresponding to January 20, 2020. The yearly average reproductive number $\bar{R}_0 = 3.2$ was supposed to be lower than in Germany due to the overall lower population density and the lower latitude (corresponding to higher UV radiation). Due to the lower latitude and higher UV radiation, seasonal fluctuations in the base reproductive number of 35% were assumed, which are lower than in Germany. The peak in $R_0$ occurred on day $t_{R_{0\max}} = 335$, which again corresponds to late December.

The general contact reductions for the USA differed substantially from those in Germany, reflecting the governments' different responses to the epidemic. The general contact reductions are described in Table S7 Table. We assumed travel and contact restrictions that reduced contacts by 55% (and hence were less restrictive than the hard lockdown in Germany) were put in place at time $t_{\mathrm{Dist}_1} = 50$ (early March), which were lifted on day $t_{\mathrm{Dist}_2} = 115$ (mid-May). After mid-May a relief period (with contact reductions of 22%) was assumed until $t_{\mathrm{Dist}_3} = 190$ (late July) after which a 'hard lockdown starts', which is as severe as the original restrictions (55%) sustained until October 1st ($t_{\mathrm{Dist}_4} = 255$), after which a moderate relief (contact reductions of 45%) occurred during the US elections until November 5 ($t_{\mathrm{Dist}_5} = 290$). After the US elections, a second "hard lockdown" (with contact reductions of 65%) was assumed until thanksgiving holidays ($t_{\mathrm{Dist}_6} = 309$). A relief period during these holidays (55% contact reduction) was sustained for 7 days (until $t_{\mathrm{Dist}_7} = 316$), after which the hard lockdown was continued with 60% contact reductions until December 10 ($t_{\mathrm{Dist}_8} = 325$), after which a severe pre-holiday-season lockdown started with contact reductions of 70% for 10 days (until $t_{\mathrm{Dist}_9} = 335$). During the holiday season from December 20 ($t_{\mathrm{Dist}_9} = 335$) until January 8, 2021 ($t_{\mathrm{Dist}_{10}} = 354$), relieved contact reductions of 55% were assumed. Finally, a hard lockdown with 65% contact reductions was assumed until mid-April 2021 ($t_{\mathrm{Dist}_{11}} = 450$). This is supposed to reflect more restrictive epidemic management under the new administration. For simplicity – and to be able to study the effect of the vaccination – after day $t_{\mathrm{Dist}_7} = 450$, all general contact reductions were lifted in the simulations.

### No vaccination

The simulations until February 2021 reflect the number of cases in the USA and hence are qualitatively different from those in Germany. After the contact reductions are lifted, the dynamics in the absence of vaccinations are qualitatively similar to those in Germany. Namely, a severe epidemic peak would emerge in fall 2021 (cf. S2 FigA-D with Figs 4A-D).

### Vaccination campaigns

Our baseline assumptions for the vaccination campaigns were an early onset of the campaign in December ($t = 330$), a vaccination coverage of 60%, an average waiting time of 180 days to get vaccinated, and a vaccination schedule of 28 days. Vaccination campaigns have qualitatively the same effect as in Germany. However, there are quantitative differences. The main difference is that vaccination coverage of 60% is sufficient to guarantee a return to normalcy – namely, only a shallow peak would emerge in spring 2022, with incidences being lower than in spring 2020. This is because of the smaller basic reproduction number that was assumed. Furthermore, the number of cases per 100 000 is higher, so that a higher level of immunity is already reached. A late start of the vaccination campaign for a broad part of the population (mid-January) would lead to a small wave in spring 2022 (if the vaccination rate would not be increased), leading to higher mortality (see S2 Fig).

Even with an early launch of the vaccination campaign ($t = 330$), vaccination coverage of 40% is too low to prevent a strong epidemic peak in spring 2022, which would be higher than the one in the holiday season 2020 (see S3 Fig). Low coverage of 25% would even result in a higher peak in fall 2021 than without vaccination. This is because the peak occurs later in the flu season due to seasonal fluctuations in $R_0$. However, the number of infections is still much lower, as the higher peak is much narrower. Coverages above 60% prevent such an epidemic peak. Importantly the waiting time to get vaccinated must be sufficiently short, to reach high levels of immunization on time. An epidemic peak, that reaches the levels of November 2020, would occur in spring 2022 if the average waiting time is 240 days (see S4 Fig). If the waiting time is even 300 days, this peak would be higher than the one experienced in the holiday season 2020 (see S4 Fig). An average waiting time of 180 days is sufficient to prevent such an outbreak (see S4 Fig).

The vaccination schedule does not have a substantial effect in the long run. However, it contributes to reducing the number of infections in the short run. A shorter vaccination cycle is advantageous (see S5 Fig) – however, with a shorter vaccination schedule, the vaccination rate might drop due to limited supply of the vaccine.

If the effectiveness of the vaccine is low (78%), a higher vaccination coverage than 60% is necessary to avoid an epidemic peak in spring 2022 – in fact, it can be avoided with a vaccination coverage of 75%. In comparison, poor effectiveness of 50% is insufficient to avoid another epidemic peak (see S6 Fig).

As for Germany ADE does not play an important role in vaccination campaigns (see S7 Fig).
