## Supplementary material for "The impact of COVID-19 vaccination campaigns accounting for antibody-dependent enhancement": Figure S2

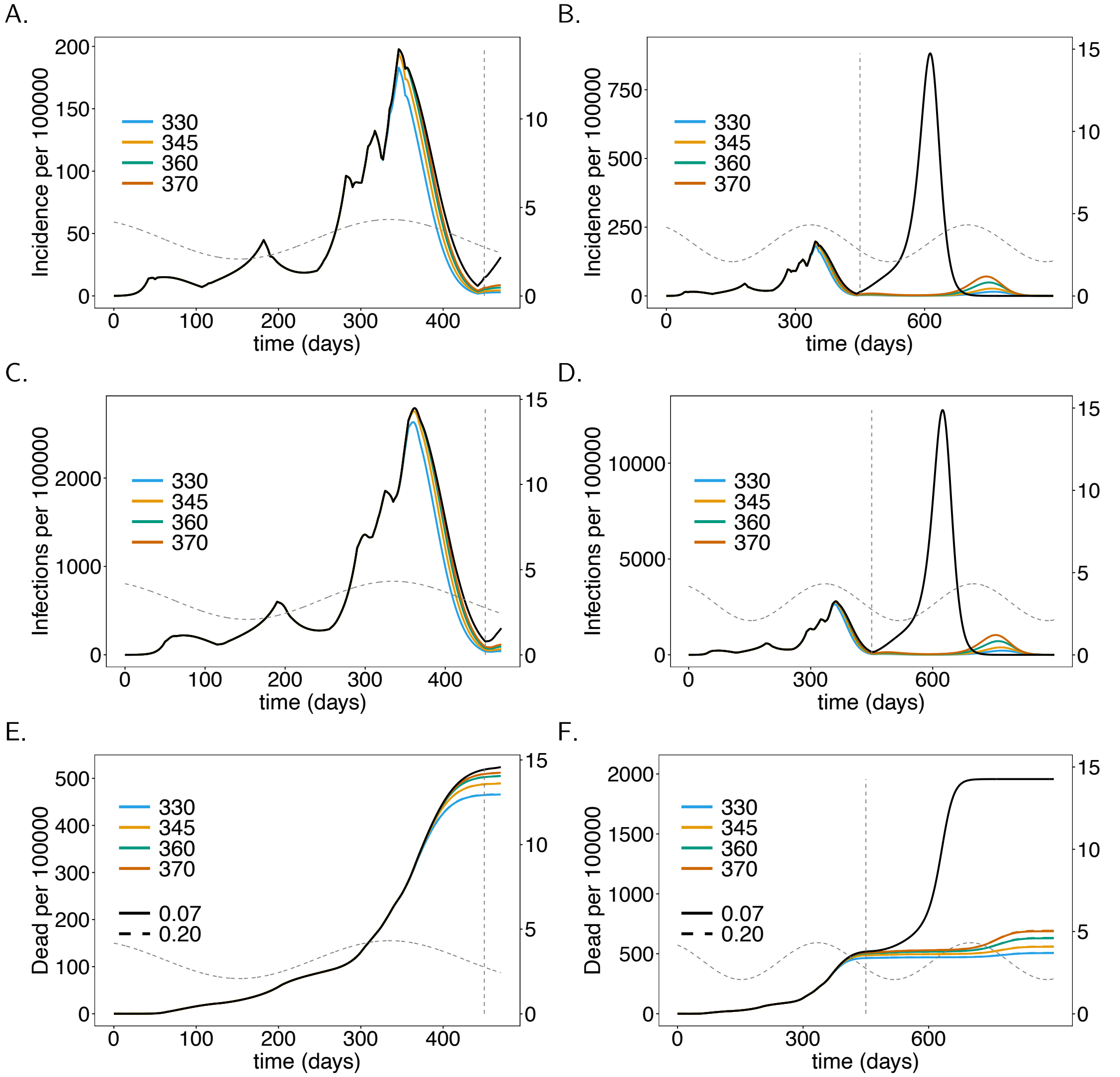


**S2 Fig. Onset of the vaccination campaign:** As in Fig 4 but for U.S. instead of Germany. Parameters for contact reduction are given in S7 Table.
