## Supplementary material for "The impact of COVID-19 vaccination campaigns accounting for antibody-dependent enhancement": Table S2

**S2 Table.** Parameters describing disease progression for Germany (GER) and the USA.

| Parameter | Description | Value |
|---|---|---|
| $n_E$ | No. of latent sub-states (Erlang states) | 16 |
| $n_P$ | No. of prodromal sub-states (Erlang states) | 16 |
| $n_I$ | No. of fully infectious sub-states (Erlang states) | 16 |
| $n_L$ | No. of late infectious sub-states (Erlang states) | 16 |
| $D_E$ | Average duration of latent period | 3.7 days |
| $D_P$ | Average duration of prodromal period | 1 day |
| $D_I$ | Average duration of fully infectious period | 5 days |
| $D_L$ | Average duration of late infectious period | 5 days |
| $\varepsilon$ | Transition rate of latent sub-states | $n_{\mathrm{E}}/D_E$ |
| $\varphi$ | Transition rate of prodromal sub-states | $n_{\mathrm{P}}/D_{\mathrm{P}}$ |
| $\gamma$ | Transition rate of fully infectious sub-states | $n_{\mathrm{I}}/D_{\mathrm{I}}$ |
| $\delta$ | Transition rate of late infectious sub-states | $n_L/D_L$ |
| $\alpha$ | Average waiting time for the outcome of the vaccine | $1/28, 1/42$ |
| $\nu$ | Rate at which individuals get vaccinated | $0, 1/180, 1/240, 1/300$ |

Parameters describing disease progression and their values used in the simulations. Abbreviations: No. . . . Number.
