## Supplementary figures and images for "The impact of COVID-19 vaccination campaigns accounting for antibody-dependent enhancement"

### Figure S1

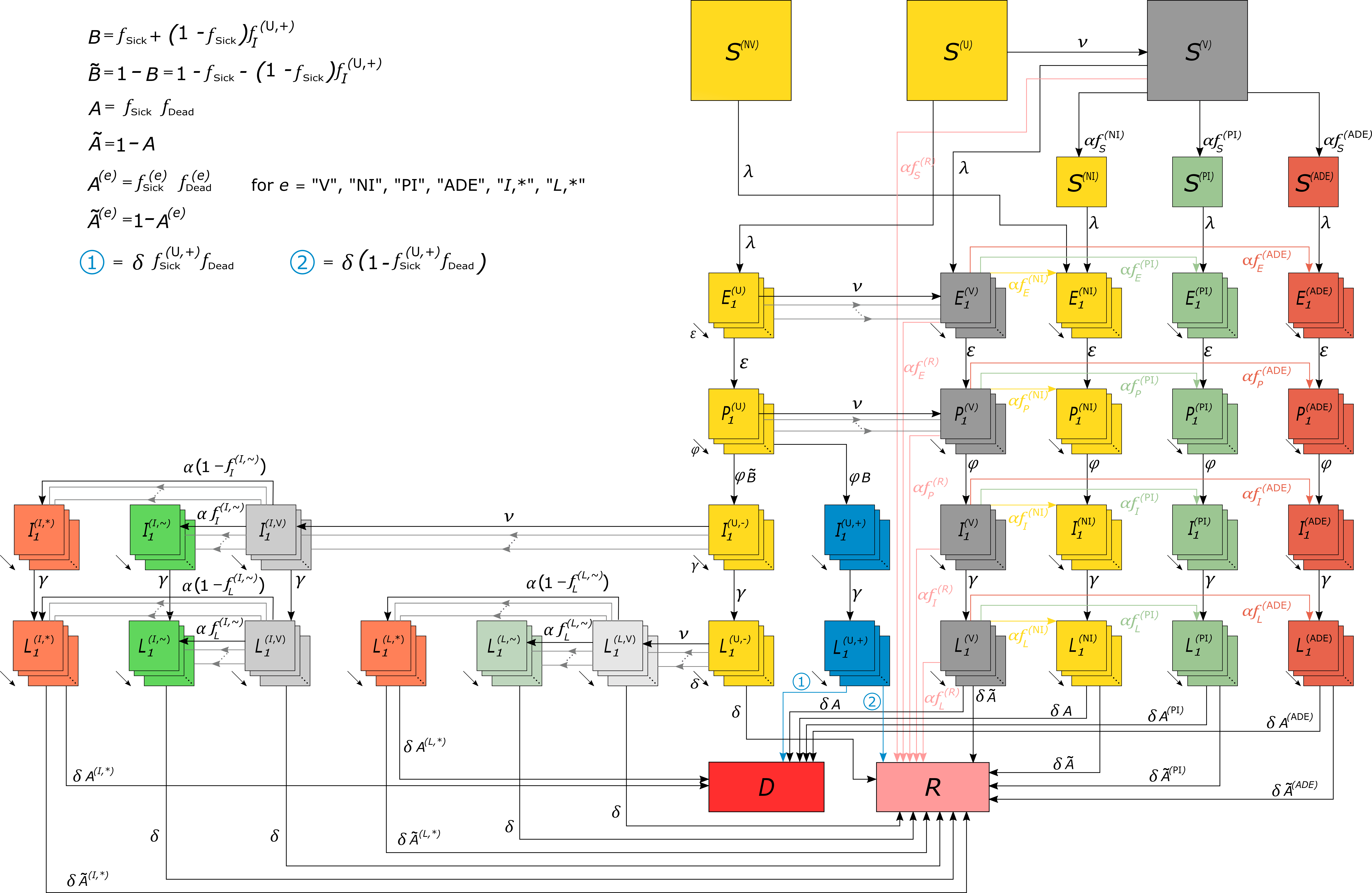

### Figure S3

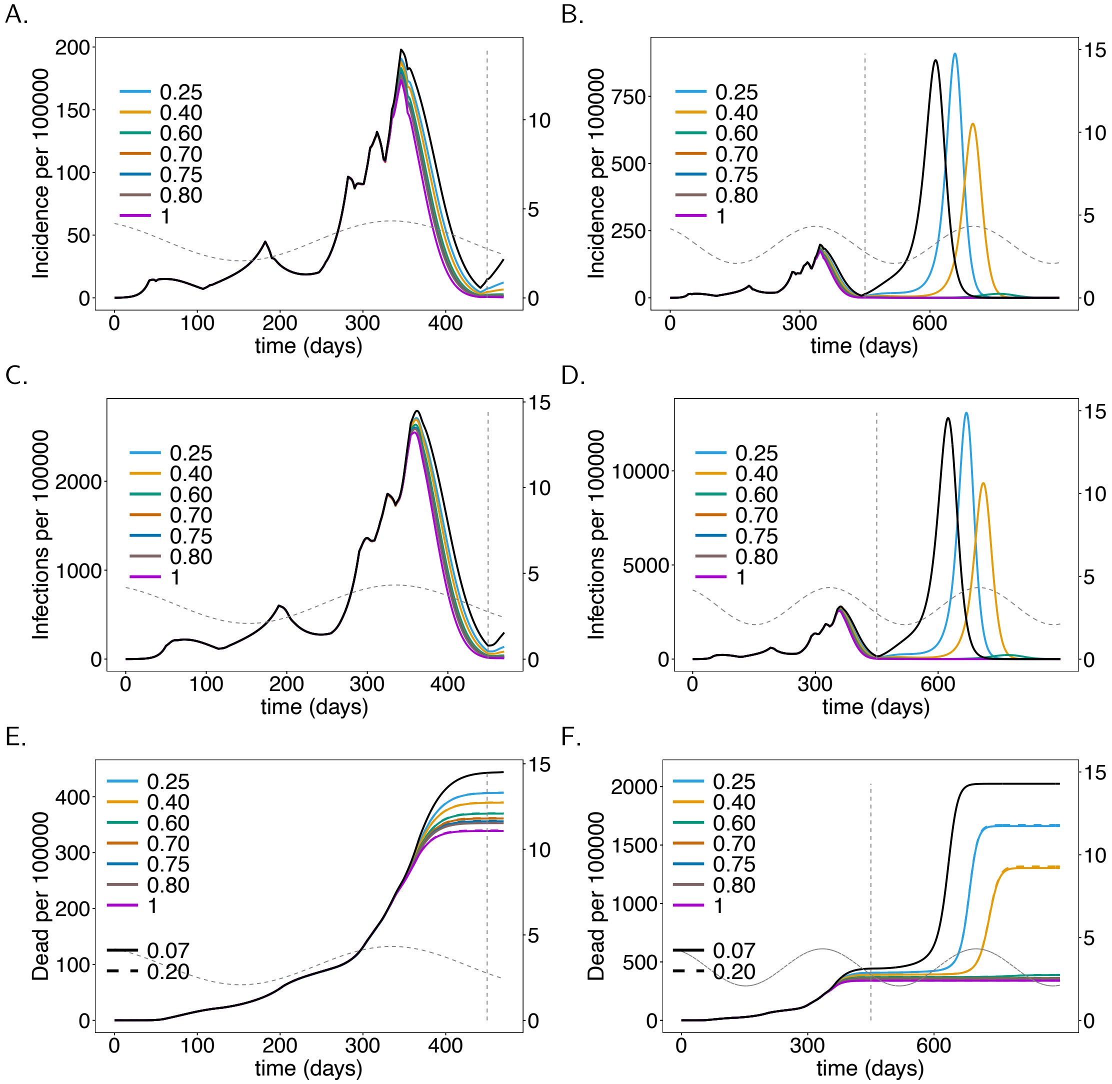


**S3 Fig. Vaccination coverage.** As Fig 5, but for the USA instead of Germany.

### Figure S4

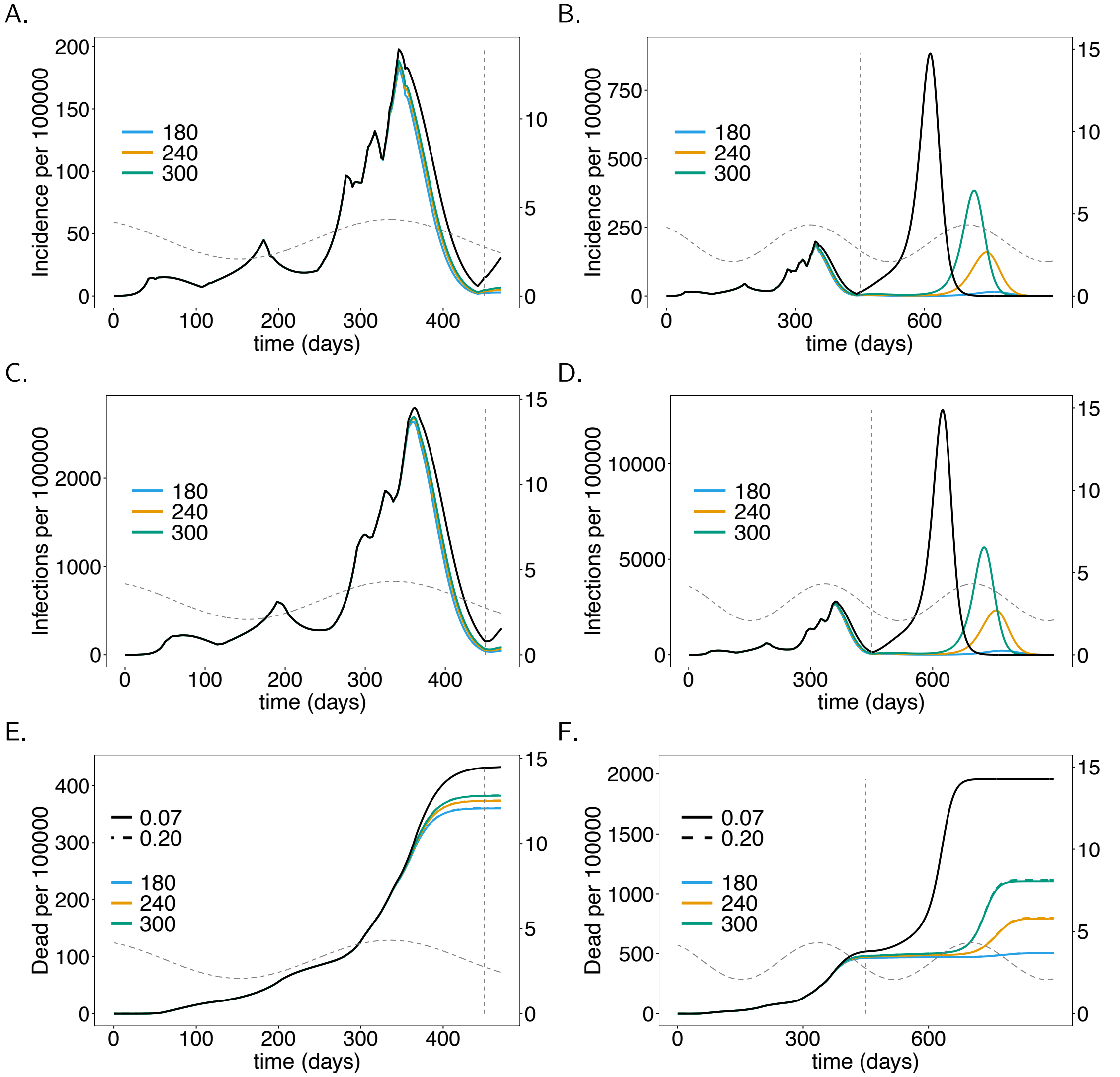


**S4 Fig. Vaccination rate.** As Fig 6, but for the USA instead of Germany.

### Figure S5

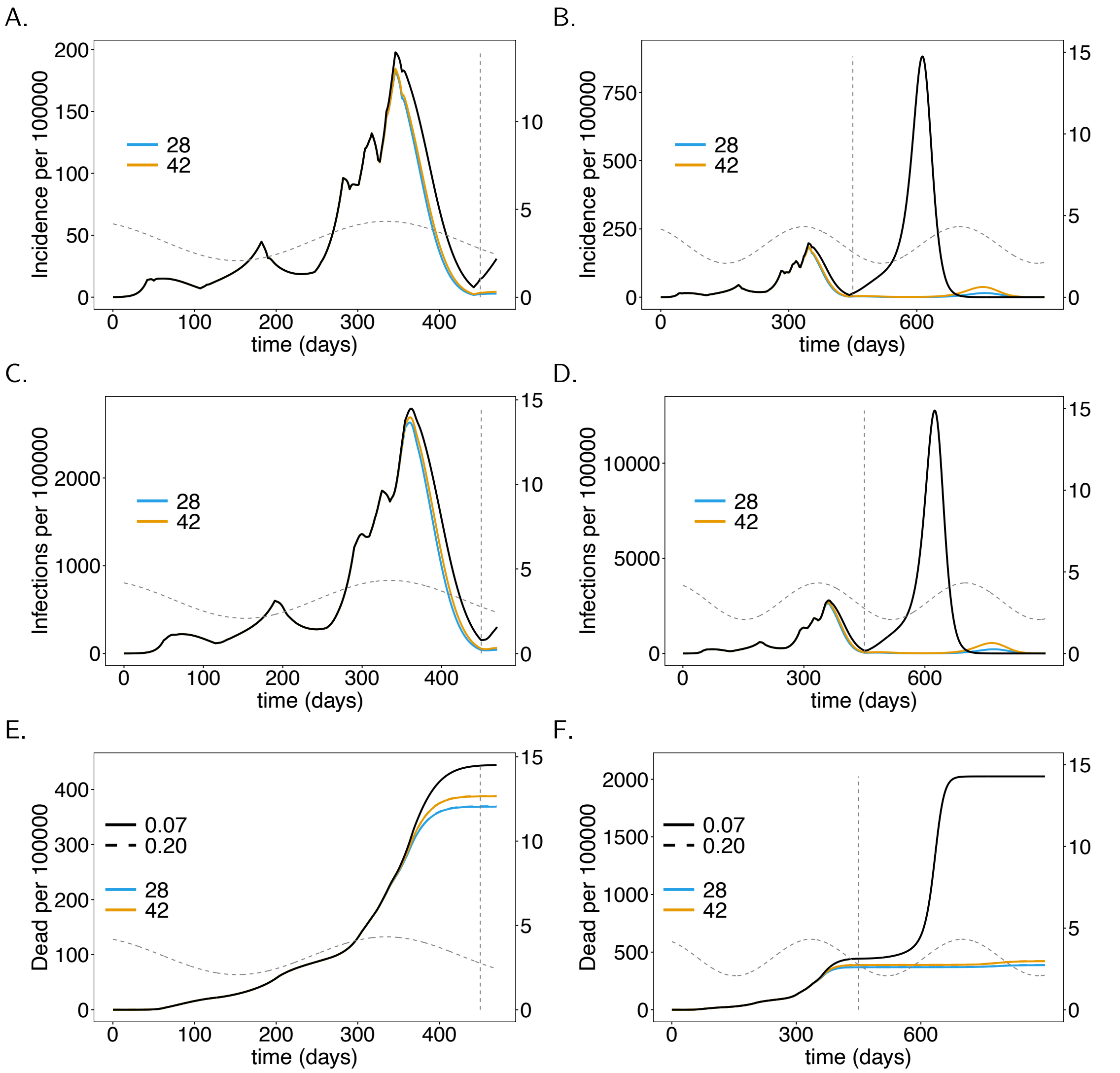


**S5 Fig. Vaccination schedule and time to immune response.** As Fig 7, but for the USA instead of Germany.

### Figure S6

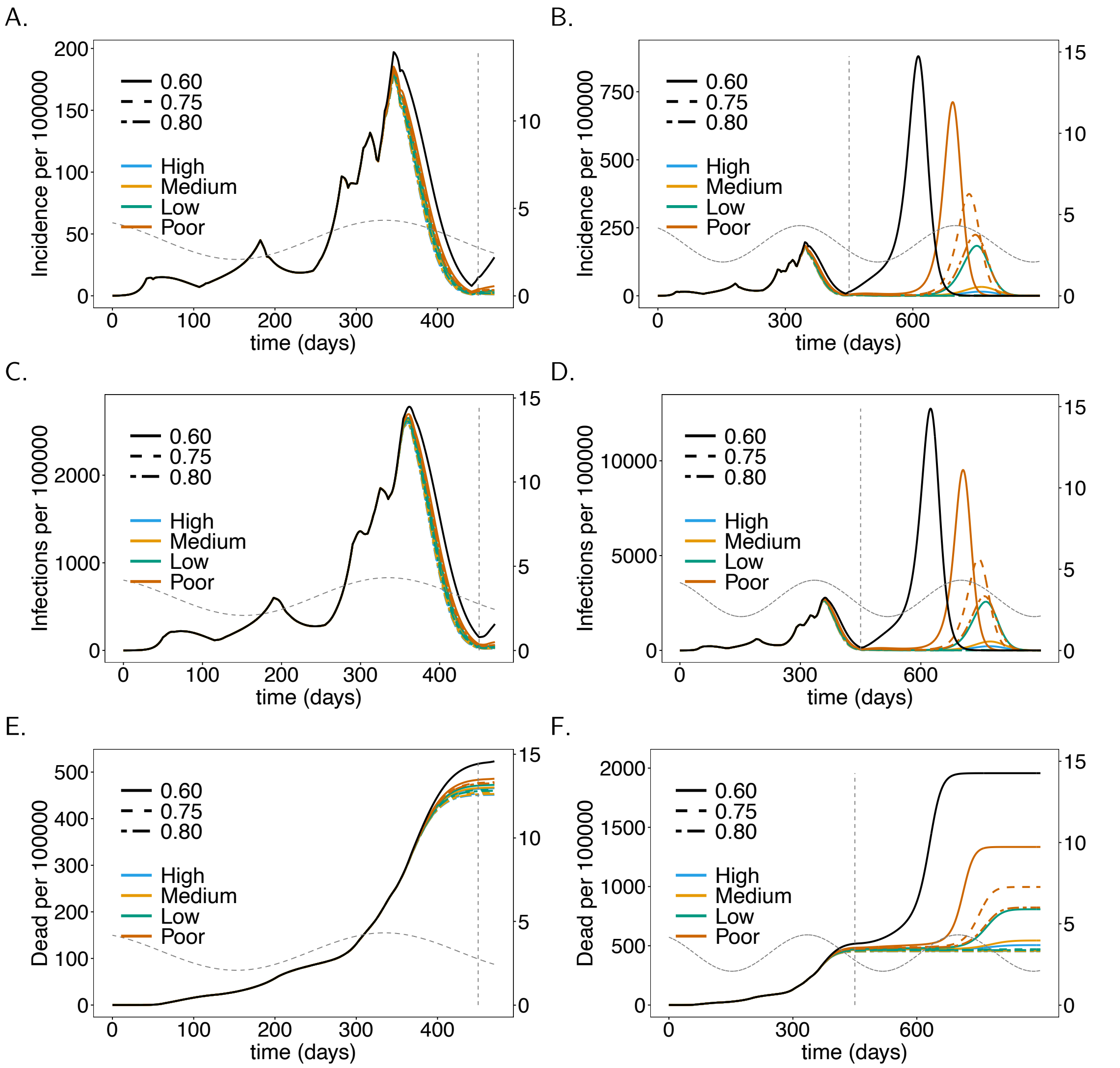


**S6 Fig. Vaccine effectiveness.** As Fig 8, but for the USA instead of Germany.

### Figure S7

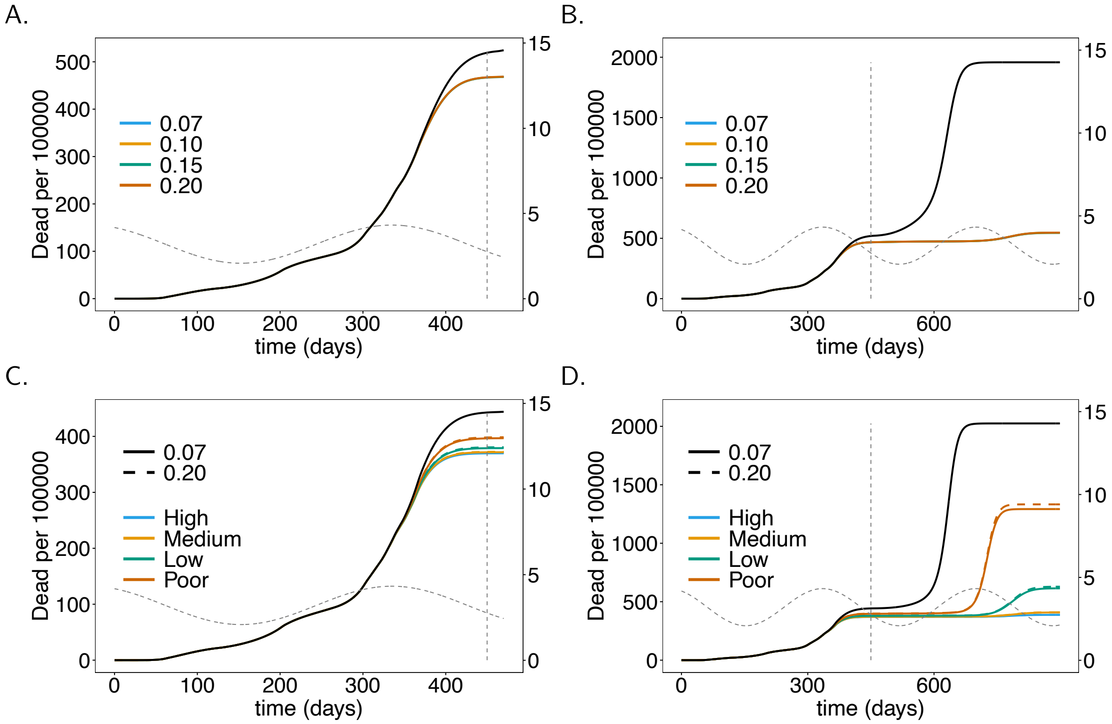


**S7 Fig. ADE-induced increased mortality.** As Fig 9, but for the USA instead of Germany.
