## Supplementary material for "The impact of COVID-19 vaccination campaigns accounting for antibody-dependent enhancement": Table S3

**S3 Table.** Summary of variables describing sub-population sizes in Germany (GER) and the USA.

| Name | Description | Initial Values | |
|---|---|---|---|
| | | GER | USA |
| $S^{(\mathrm{NV})}(t)$ | Unvaccinable susceptibles (anti-vaxxers & inds. that cannot be vaccinated) | 33 200 000 | 132 400 000 |
| $S^{(\mathrm{U})}(t)$ | Susceptibles waiting to be vaccinated | 49 799 800 | 198 599 925 |
| $S^{(\mathrm{V})}(t)$ | Vaccinated susceptibles with pending vaccine outcome | | 0 |
| $S^{(\mathrm{NI})}(t)$ | Vaccinated susceptibles that failed to immunize | | 0 |
| $S^{(\mathrm{PI})}(t)$ | Vaccinated susceptibles who developed partial immunity | | 0 |
| $S^{(\mathrm{ADE})}(t)$ | Vaccinated susceptibles who developed ADE | | 0 |
| $E_k^{(\mathrm{U})}(t)$ | Latently inf. inds. waiting to be vaccinated | | 0 |
| $E_k^{(\mathrm{V})}(t)$ | Vaccinated latently inf. inds. with pending vaccine outcome | | 0 |
| $E_k^{(\mathrm{NI})}(t)$ | Unvaccinable latently inf. inds. and vaccinated ones who failed to immunize | | 0 |
| $E_k^{(\mathrm{PI})}(t)$ | Vaccinated latently inf. inds. who developed partial immunity | | 0 |
| $E_k^{(\mathrm{ADE})}(t)$ | Vaccinated latently inf. inds. who developed ADE | | 0 |
| $P_k^{(\mathrm{U})}(t)$ | Vaccinated prodromal inds. waiting to be vaccinated | | 0 |
| $P_k^{(\mathrm{V})}(t)$ | Vaccinated prodromal inds. with pending vaccine outcome | | 0 |
| $P_k^{(\mathrm{NI})}(t)$ | Unvaccinable prodromal inds. and vaccinated ones who failed to immunize | | 0 |
| $P_k^{(\mathrm{PI})}(t)$ | Vaccinated prodromal inds. who developed partial immunity | | 0 |
| $P_k^{(\mathrm{ADE})}(t)$ | Vaccinated prodromal inds. who developed ADE | | 0 |
| | | GER | USA |
| $I_k^{(\mathrm{U},-)}(t)$ | Undiagnosed and asymptomatic fully inf. inds. waiting to be vaccinated | $k = 1$: 200 | 75 |
| | | $k > 1$: 0 | 0 |
| $I_k^{(\mathrm{U},+)}(t)$ | Diagnosed or symptomatic fully inf. inds. that were waiting to get vaccinated | | 0 |
| $I_k^{(\mathrm{V})}(t)$ | Fully inf. inds. with pending vaccine outcome | | 0 |
| $I_k^{(\mathrm{NI})}(t)$ | Unvaccinable fully inf. inds. and vaccinated ones who failed to immunize | | 0 |
| $I_k^{(\mathrm{PI})}(t)$ | Vaccinated fully inf. inds. who developed partial immunity | | 0 |
| $I_k^{(\mathrm{ADE})}(t)$ | Vaccinated fully inf. inds. who developed ADE | | 0 |
| $I_k^{(I,\mathrm{V})}(t)$ | Fully inf. inds. who were vaccinated during this phase with pending vaccine outcome | | 0 |
| $I_k^{(I,\sim)}$ | Fully inf. inds. who were vaccinated during this phase and had a neutral outcome | | 0 |
| $I_k^{(I,*)}$ | Fully inf. inds. who were vaccinated during this phase and had a del. outcome | | 0 |
| $L_k^{(\mathrm{U},+)}(t)$ | Diagnosed or symptomatic late inf. inds. that were waiting to get vaccinated | | 0 |
| $L_k^{(\mathrm{U},-)}(t)$ | Undiagnosed and asymptomatic late inf. inds. waiting to be vaccinated | | 0 |
| $L_k^{(\mathrm{V})}(t)$ | Late inf. inds. with pending vaccine outcome | | 0 |
| $L_k^{(\mathrm{NI})}(t)$ | Unvaccinable late inf. inds. and vaccinated ones who failed to immunize | | 0 |
| $L_k^{(\mathrm{PI})}(t)$ | Vaccinated late inf. inds. who developed partial immunity | | 0 |
| $L_k^{(\mathrm{ADE})}(t)$ | Vaccinated late inf. inds. who developed ADE | | 0 |
| $L_k^{(I,\mathrm{V})}(t)$ | Late inf. inds. who were vaccinated while fully infectious with pending vaccine outcome | | 0 |
| $L_k^{(I,\sim)}(t)$ | Late inf. inds. who were vaccinated while fully infectious and had a neutral outcome | | 0 |
| $L_k^{(I,*)}(t)$ | Late inf. inds. who were vaccinated while fully infectious and had a del. outcome | | 0 |
| $L_k^{(L,\mathrm{V})}(t)$ | Late inf. inds. who were vaccinated during this phase with pending vaccine outcome | | 0 |
| $L_k^{(L,\sim)}(t)$ | Late inf. inds. who were vaccinated during this phase and had a neutral outcome | | 0 |
| $L_k^{(L,*)}(t)$ | Late inf. inds. who were vaccinated during this phase and had a del. outcome | | 0 |
| $R(t)$ | Recovered or fully immunized individuals | | 0 |
| $D(t)$ | Dead individuals | | 0 |

Summary of all model compartments and their initial values chosen for simulation. Abbreviations: del.. . . deleterious; inf. . . . infectious; inds. . . . individuals.
