## Supplementary material for "The impact of COVID-19 vaccination campaigns accounting for antibody-dependent enhancement": Table S4

**S4 Table.** Parameters describing disease severity and mortality for Germany (GER) and the USA.

| Parameters | Description | | Value |
|---|---|---|---|
| | Fraction of ... | | |
| $f_S^{(\mathrm{NI})}$ | ... susceptibles that fails to immunize after vaccination | | cf. Table 1 |
| $f_S^{(\mathrm{PI})}$ | ... susceptibles that develops partial immunity after vaccination | | - |
| $f_S^{(\mathrm{ADE})}$ | ... susceptibles that develops ADE after vaccination | | - |
| $f_S^{(R)}$ | ... susceptibles that develops permanent immunity after vaccination | | - |
| $f_E^{(\mathrm{NI})}$ | ... latently inf. that fails to immunize after vaccination | | cf. Table 1 |
| $f_E^{(\mathrm{PI})}$ | ... latently inf. inds. that develops partial immunity after vaccination | | - |
| $f_E^{(\mathrm{ADE})}$ | ... latently inf. that develops ADE after vaccination | | - |
| $f_E^{(R)}$ | ... latently inf. that develops permanent immunity | | - |
| $f_P^{(\mathrm{NI})}$ | ... prodromal inds. that fails to immunize after vaccination | | cf. Table 1 |
| $f_P^{(\mathrm{PI})}$ | ... prodromal inds. that develops partial immunity after vaccination | | - |
| $f_P^{(\mathrm{ADE})}$ | ... prodromal inds. that develops ADE after vaccination | | - |
| $f_P^{(R)}$ | ... prodromal inds. that develops permanent immunity after vaccination | | - |
| $f_I^{(\mathrm{U},+)}$ | ... detected or sympt. fully inf. inds. that were waiting for vaccination | | 0.005 |
| $f_I^{(\mathrm{NI})}$ | ... fully inf. inds. that fails to immunize after vaccination | | cf. Table 1 |
| $f_I^{(\mathrm{PI})}$ | ... fully inf. inds. that develops partial immunity after vaccination | | - |
| $f_I^{(\mathrm{ADE})}$ | ... fully inf. inds. that develops ADE after vaccination | | - |
| $f_I^{(R)}$ | ... fully inf. inds. that develops permanent immunity after vaccination | | - |
| $f_I^{(I,\sim)}$ | ... fully infectious inds. vaccinated during this phase with neutral outcome | | 0.95 |
| $f_L^{(\mathrm{NI})}$ | ... late inf. inds. that fails to immunize after vaccination | | 0 |
| $f_L^{(\mathrm{PI})}$ | ... late inf. inds. that develops partial immunity after vaccination | | 0 |
| $f_L^{(\mathrm{ADE})}$ | ... late inf. inds. that develops ADE after vaccination | | 0 |
| $f_L^{(R)}$ | ... late inf. inds. that develops permanent immunity after vaccination | | 0 |
| $f_L^{(I,\sim)}$ | ... late infectious inds. vaccinated during the fully inf. phase with neutral outcome | | 0.95 |
| $f_L^{(L,\sim)}$ | ... late infectious inds. vaccinated during this phase with neutral outcome | | 0.99 |
| | | GER | USA |
| $f_{\mathrm{Sick}}$ | ... sympt. individuals | 0.58 | 0.58 |
| $f_{\mathrm{Dead}}$ | ... sympt. inds. who die from COVID-19 | 0.016 | 0.04 |
| $f_{\mathrm{Sick}}^{(\mathrm{PI})}$ | ... sympt. inds. with partial immunity | | 0.50 |
| $f_{\mathrm{Dead}}^{(\mathrm{PI})}$ | ... sympt. inds. with partial immunity who die from COVID-19 | | 0.02 |
| $f_{\mathrm{Dead}}^{(\mathrm{ADE})}$ | ... sympt. inds. with ADE who die from COVID-19 | | 0.07, 0.2 |
| $f_{\mathrm{Sick}}^{(\mathrm{U},+)}$ | ... sympt. or diagnosed individuals that were waiting to be vaccinated | | cf. eq. 3 |
| $f_{\mathrm{Sick}}^{(\mathrm{ADE})}$ | ... sympt. inds. with ADE | | 0.92 |
| $f_{\mathrm{Sick}}^{(I,*)}$ | ... sympt. fully inf. inds. vaccinated during this phase with del. outcome | | 0.58 |
| $f_{\mathrm{Dead}}^{(I,*)}$ | ... lethal infs. among fully inf. inds. vaccinated during this phase with del. outcome | | 0.07 |
| $f_{\mathrm{Sick}}^{(L,*)}$ | ... sympt. late inf. inds. vaccinated during this phase with del. outcome | | 0.58 |
| $f_{\mathrm{Dead}}^{(L,*)}$ | ... lethal infs. among late inf. inds. vaccinated during this phase with del. outcome | | 0.07 |

Abbreviations: del. ... deleterious; eq. ... Equation; inf. ... infectious; infs. ... infections; inds. ... individuals; sympt. ... symptomatic.
