## Supplementary material for "The impact of COVID-19 vaccination campaigns accounting for antibody-dependent enhancement": Table S5

**S5 Table.** Parameters describing contact behavior and force of infection for Germany (GER) and the USA.

| Parameter | Definition | Value/Eq. | |
|---|---|---|---|
| | | GER | USA |
| $\lambda_{\text{Ext}}$ | Infections from outside of the population | 45/day | 50/day |
| $\bar{R}_0$ | Annual average basic reproduction number | 3.4 | 3.2 |
| $a$ | Amplitude of the seasonal fluctuation in $R_0$ | 0.43 | 0.35 |
| $t_{R_{0\max}}$ | Day when $R_0$ reaches its maximum | 300 | 335 |
| $Q_{\max}$ | Maximum capacity of isolation units per 10,000 | 200 | 30 |
| $t_{\text{Iso}_1}$ | Day case isolation measures start | 30 | 20 |
| $t_{\text{Iso}_2}$ | Day case isolation measures end | 900 | 900 |
| $p_{\text{Home}}$ | Contact reduction in home isolation | | 75% |
| $c_P$ | Relative contagiousness in prodromal period | | 0.5 |
| $c_I$ | Relative contagiousness in fully inf. phase | | 1 |
| $c_L$ | Relative contagiousness in late inf. phase | | 0.5 |
| $\beta_P(t)$ | Seasonally varying effective contact rate of prodromal inds. | | cf. eq. 7a |
| $\beta_I(t)$ | Seasonally varying effective contact rate of fully infectious inds. | | cf. eq. 7b |
| $\beta_L(t)$ | Seasonally varying effective contact rate of late infectious inds. | | cf. eq. 7c |

Abbreviations: eq. …Equation; inf. …infectious; inds. …individuals.
