## Supplementary material for "The impact of COVID-19 vaccination campaigns accounting for antibody-dependent enhancement": Table S6

**S6 Table.** Contact reduction parameters chosen for the simulations of Germany.

| Parameter | Description | Value |
|---|---|---|
| $t_{\mathrm{Dist}_1}$ | Day when first "hard lockdown" (general distancing) starts | 40 |
| $t_{\mathrm{Dist}_2}$ | Day when first "hard lockdown" ends and first "relief period" starts | 82 |
| $t_{\mathrm{Dist}_3}$ | Day when first "relief period" ends and "soft lockdown" starts | 246 |
| $t_{\mathrm{Dist}_4}$ | Day when "soft lockdown" ends and the second "hard lockdown" starts | 280 |
| $t_{\mathrm{Dist}_5}$ | Day when second "hard lockdown" ends and relief period starts | 397 |
| $t_{\mathrm{Dist}_6}$ | Day when general contact reduction ends | 450 |
| $p_{\mathrm{Cont}_1}$ | General contact reduction between individuals during the first lockdown | 70% |
| $p_{\mathrm{Cont}_2}$ | General contact reduction during the first "relief period" | 40% |
| $p_{\mathrm{Cont}_3}$ | General contact reduction during the "soft lockdown" | 50% |
| $p_{\mathrm{Cont}_4}$ | General contact reduction between individuals during the "hard lockdown" | 68% |
| $p_{\mathrm{Cont}_5}$ | General contact reduction between individuals during the second "relief period" | 50% |
