## Supplementary material for "The impact of COVID-19 vaccination campaigns accounting for antibody-dependent enhancement": Table S7

**S7 Table.** Contact reduction parameters chosen for the simulations of the USA.

| Parameter | Description | Value |
|---|---|---|
| $t_{\mathrm{Dist}_1}$ | Day when first contact restrictions (e.g. travel bans etc.) start | 50 |
| $t_{\mathrm{Dist}_2}$ | Day when restrictions were partly lifted | 115 |
| $t_{\mathrm{Dist}_3}$ | Day when first “relief period” ends and a “hard lockdown” starts | 190 |
| $t_{\mathrm{Dist}_4}$ | Day when “hard lockdown” is relieved into a “soft lockdown” | 255 |
| $t_{\mathrm{Dist}_5}$ | Day when second “soft lockdown” ends and a new “hard lockdown” starts | 290 |
| $t_{\mathrm{Dist}_6}$ | Day when second “hard lockdown” ends and a “relief period” starts | 309 |
| $t_{\mathrm{Dist}_7}$ | Day when “relief period” ends and a subsequent “hard lockdown” starts | 316 |
| $t_{\mathrm{Dist}_8}$ | Day when “hard lockdown” ends and a stronger lockdown starts | 325 |
| $t_{\mathrm{Dist}_9}$ | Day when stronger “hard lockdown” ends and a “soft lockdown” starts | 335 |
| $t_{\mathrm{Dist}_{10}}$ | Day when “soft lockdown” ends and “hard lockdown” starts | 354 |
| $t_{\mathrm{Dist}_{11}}$ | Day when general contact reduction ends | 450 |
| $p_{\mathrm{Cont}_1}$ | General contact reduction between individuals during the first lockdown | 55% |
| $p_{\mathrm{Cont}_2}$ | General contact reduction during the first “relief period” | 22% |
| $p_{\mathrm{Cont}_3}$ | General contact reduction during the “hard lockdown” | 55% |
| $p_{\mathrm{Cont}_4}$ | General contact reduction between individuals during the “soft lockdown” | 45% |
| $p_{\mathrm{Cont}_5}$ | General contact reduction between individuals during the second “hard lockdown” | 65% |
| $p_{\mathrm{Cont}_6}$ | General contact reduction between individuals during the second “relief period” | 55% |
| $p_{\mathrm{Cont}_7}$ | General contact reduction during the “hard lockdown” | 60% |
| $p_{\mathrm{Cont}_8}$ | General contact reduction during the stronger “hard lockdown” | 70% |
| $p_{\mathrm{Cont}_9}$ | General contact reduction between individuals during the “soft lockdown” | 55% |
| $p_{\mathrm{Cont}_{10}}$ | General contact reduction between individuals during the “hard lockdown” | 65% |
