## Supplementary material for "The impact of COVID-19 vaccination campaigns accounting for antibody-dependent enhancement": S1 Mathematical Appendix

### Appendix - Mathematical description

Here we concisely formulate the mathematical model for the occurrence of ADE during vaccination campaigns. We generalize the SEIR model underlying the pandemic-preparedness-tool CovidSim (cf. [1]). The presentation follows [2] and [3], two very different generalizations of the CovidSim 1.1 model tailored to study the spread of COVID-19 in closed facilities and multiple exposures to the virus, respectively. To adequately model the occurrence of ADE during vaccination campaigns, the population needs to be partitioned into several groups, and hence into a large number of compartments in the SEIR model. We therefore, describe the model in a step-wise procedure. Figures 1 - 3 are simplified illustrations intended to facilitate the understanding of the model's complex flow chart (S1 Fig). In the following, regarding the model's compartments, we use superscripts, to refer to sub-divided groups. If we are not referring to a particular sub-group we use a (.) as a placeholder.

#### Model compartments

In a population of size $N$, the model follows the numbers of individuals being susceptible ($S^{(.)}$), in the latent phase ($E^{(.)}_{\text{Sum}}$) consisting of $n_E$ equivalent sub-states ($E^{(.)}_k$, $k = 1, \ldots, n_E$), in the prodromal phase ($P^{(.)}_{\text{Sum}}$) consisting of $n_P$ equivalent sub-states ($P^{(.)}_k$, $k = 1, \ldots, n_P$), the fully infectious phase ($I^{(.)}_{\text{Sum}}$) consisting of $n_I$ equivalent sub-states ($I^{(.)}_k$, $k = 1, \ldots, n_I$), the late infectious phase ($L^{(.)}_{\text{Sum}}$) consisting of $n_L$ equivalent sub-states ($L^{(.)}_k$, $k = 1, \ldots, n_L$), the final "removed" state ($R$), and dead individuals ($D$). Therefore, the number of individuals in the latent, prodromal, fully infectious and late infectious phases in the respective sub-groups are given by

$$E^{(.)}_{\text{Sum}}(t) = \sum_{k=1}^{n_E} E^{(.)}_k(t), \tag{1a}$$

$$P^{(.)}_{\text{Sum}}(t) = \sum_{k=1}^{n_P} P^{(.)}_k(t), \tag{1b}$$

$$I^{(.)}_{\text{Sum}}(t) = \sum_{k=1}^{n_I} I^{(.)}_k(t), \tag{1c}$$

and

$$L^{(.)}_{\text{Sum}}(t) = \sum_{k=1}^{n_L} L^{(.)}_k(t). \tag{1d}$$

The sub-states are used to avoid naive exponentially-distributed times during disease progression, which implicitly occur in SEIR models. By the equivalent sub-states the waiting times become Erlang-distributed, i.e., the variance of the durations are not determined by their mean [1, 2].

#### Susceptible individuals

Susceptible individuals are sub-divided (see Fig 2) into those that are: (i) not vaccinable because they do not have access, refuse to be vaccinated, or cannot be vaccinated for medical reasons (e.g., allergies, or the vaccine is not approved for them), denoted by ($S^{(\mathrm{NV})}$); (ii) waiting to be vaccinated ($S^{(\mathrm{U})}$); (iii) already vaccinated, but the outcome of the vaccination is still pending ($S^{(\mathrm{V})}$); (iv) already vaccinated but only partially immunized ($S^{(\mathrm{PI})}$); (v) vaccinated, but the vaccination failed to immunize at all ($S^{(\mathrm{NI})}$); (vi) vaccinated, but the vaccine caused ADE ($S^{(\mathrm{ADE})}$). The waiting time for susceptibles ($S^{(\mathrm{U})}$) to get vaccinated is $D_V$. The effect of the vaccine is not immediate (pending immunization). Thus, individuals from ($S^{(\mathrm{U})}$), if not infected, are moved into compartment ($S^{(\mathrm{V})}$) at rate $\nu = 1/D_V$. If not infected, after an average duration $D_A$, the outcome of the vaccine manifests and these susceptibles either are (i) fully immune ($R$) with probability $f_S^{(R)}$; (ii) partially immune ($S^{(\mathrm{PI})}$, i.e., they can still get infected) with probability $f_S^{(\mathrm{PI})}$; (iii) not immune ($S^{(\mathrm{NI})}$, i.e., the vaccine had no effect) with probability $f_S^{(\mathrm{NI})}$; or (iv) develop ADE ($S^{(\mathrm{ADE})}$, i.e., the vaccine had a deleterious effect) with probability $f_S^{(\mathrm{ADE})}$. Hence, individuals (if not infected) are moved from $S^{(\mathrm{V})}$ to $R$, $S^{(\mathrm{PI})}$, $S^{(\mathrm{NI})}$, and $S^{(\mathrm{ADE})}$ at rates $\alpha f_S^{(R)}$, $\alpha f_S^{(\mathrm{PI})}$, $\alpha f_S^{(\mathrm{NI})}$, and $\alpha f_S^{(\mathrm{ADE})}$, respectively, where $\alpha = 1/D_A$. (Clearly, $f_S^{(R)} + f_S^{(\mathrm{PI})} + f_S^{(\mathrm{NI})} + f_S^{(\mathrm{ADE})} = 1$.)

Susceptibles that are either unvaccinable ($S^{(\mathrm{NV})}$), waiting to be vaccinated ($S^{(\mathrm{U})}$), vaccinated but have pending immunity ($S^{(\mathrm{V})}$), or failed to immunize ($S^{(\mathrm{NI})}$) are equally susceptible. Susceptibles that developed partial immunity ($S^{(\mathrm{PI})}$) or ADE ($S^{(\mathrm{ADE})}$) are less likely to become infected than the other susceptibles. Namely, only fractions $p_{\mathrm{PI}}$ and $p_{\mathrm{ADE}}$ of contacts that would infect other susceptibles are infective. If infected, partial immunity manifests in a higher likelihood of asymptomatic infections and lower mortality of symptomatic infections, while ADE results in the opposite.

#### Course of the infection

Once infected the average duration of the latent, prodromal, fully infectious, and late infectious periods are $D_E$, $D_P$, $D_I$, and $D_L$, respectively. In each of these phases individuals transit through a number of equivalent sub-stages. The reason is to avoid naive exponentially distributed durations (see above; cf. [1]). Individuals leave each latent sub-state at rate

$$\varepsilon = \frac{n_E}{D_E}. \tag{2a}$$

Similarly, the prodromal, fully infectious, and late infectious sub-states are left, respectively, at rates

$$\varphi = \frac{n_P}{D_P}, \gamma = \frac{n_I}{D_I}, \text{ and } \delta = \frac{n_L}{D_L}. \tag{2b}$$

During the latent phase, infected individuals are not yet infectious or show symptoms (see Fig 1). In the prodromal phase individuals are already infectious, however not to the full extent, and still do not show symptoms. At the beginning of the fully infectious period it is determined, whether an infection remains asymptomatic or becomes symptomatic. A fraction $f_{\text{Sick}}^{(.)}$ of fully infectious individuals becomes symptomatic, i.e., they become sick. This fraction depends on the immune status of the infected individuals, i.e., if they are partially immunized ($f_{\text{Sick}}^{(\text{PI})}$), not immunized ($f_{\text{Sick}}^{(\text{NI})}$), or developed ADE ($f_{\text{Sick}}^{(\text{ADE})}$). In particular, partially immunized individuals are less likely to develop symptoms, while individuals with ADE are more likely to develop symptoms, i.e., $f_{\text{Sick}}^{(\text{PI})} \leq f_{\text{Sick}}^{(\text{NI})} \leq f_{\text{Sick}}^{(\text{ADE})}$.

Importantly, in the fully and late infectious phases, asymptomatic infections, can turn into symptomatic infections, either when the effect of the vaccine (if still pending) manifests, or when individuals become vaccinated, potentially resulting in additional stress on the immune system. Conversely, symptomatic infections might become asymptomatic when the effect of the vaccine manifests.

Notably, vaccinated individuals ($S^{(\text{V})}$) can become infected before the effect of the immunization manifests. Then, the effect of the vaccine (complete immunity, partial immunity, no effect, or ADE) is determined during the infection. The likelihood of the three outcomes depends then on the phase of the infection. Furthermore, unvaccinated individuals can get vaccinated during an infection. Importantly, only asymptomatic individuals will be vaccinated. Particular regard is given to individuals that are vaccinated in the fully infectious and late infectious phases (see below).

##### Infection in individuals that are unvaccinable or failed to immunize

Those individuals that are unvaccinable and those that failed to immunize after being vaccinated are essentially identical. It is important to account for them separately as long as they are susceptible (cf. S1 Fig). Once they are infected, they can be modelled by the same compartments. Hence, upon infection, unvaccinable susceptibles and those that failed to immunize ($S^{(\text{NV})}$, $S^{(\text{NI})}$) are moved in the compartment $E_1^{(\text{NI})}$. From there they progress through the sub-states $E_k^{(\text{NI})}$, $P_k^{(\text{NI})}$, $I_k^{(\text{NI})}$, $L_k^{(\text{NI})}$ and finally to $R$ or $D$ (see S1 Fig, yellow compartments on the right). More precisely, a fraction $f_{\text{Sick}} f_{\text{Dead}}$ of individuals die, whereas the remaining fraction $(1 - f_{\text{Sick}} f_{\text{Dead}})$ recovers.

##### Individuals with partial immunity or ADE upon infection

Upon infection, susceptibles that developed partial immunity ($S^{(\text{PI})}$) are moved into the compartment $E_1^{(\text{PI})}$. From there they pass through the sub-states $E_k^{(\text{PI})}$, $P_k^{(\text{PI})}$, $I_k^{(\text{PI})}$, $L_k^{(\text{PI})}$ and finally a fraction $1 - f_{\text{Sick}}^{(\text{PI})} f_{\text{Dead}}^{(\text{PI})}$ recovers ($R$), whereas the remaining fraction $f_{\text{Sick}}^{(\text{PI})} f_{\text{Dead}}^{(\text{PI})}$ dies ($D$) (see S1 Fig, green compartments on the right). Likewise, susceptible individuals with ADE ($S^{(\text{ADE})}$), progress through $E_k^{(\text{ADE})}$, $P_k^{(\text{ADE})}$, $I_k^{(\text{ADE})}$, $L_k^{(\text{ADE})}$ and finally to $R$ (a fraction $1 - f_{\text{Sick}}^{(\text{ADE})} f_{\text{Dead}}^{(\text{ADE})}$) or $D$ (a fraction $f_{\text{Sick}}^{(\text{ADE})} f_{\text{Dead}}^{(\text{ADE})}$) – see S1 Fig (red compartments on the right).

##### Vaccinated individuals that get infected while the immune response is still pending

Vaccinated susceptibles for whom the effect of the vaccine is still pending upon infection ($S^{(\text{V})}$) are first moved in the compartment $E_1^{(\text{V})}$. From there, they can recover or die before or after the vaccine had an effect. In the first case (see S1 Fig, grey compartments in the middle), they progress through the states $E_k^{(\mathrm{V})}$, $P_k^{(\mathrm{V})}$, $I_k^{(\mathrm{V})}$, $L_k^{(\mathrm{V})}$, and finally $R$ or $D$. The manifestation of the infections in the compartments $E_k^{(\mathrm{V})}$, $P_k^{(\mathrm{V})}$, $I_k^{(\mathrm{V})}$, $L_k^{(\mathrm{V})}$, is identical to that in the compartments $E_k^{(\mathrm{NI})}$, $P_k^{(\mathrm{NI})}$, $I_k^{(\mathrm{NI})}$, $L_k^{(\mathrm{NI})}$, since the vaccine had no effect. In the second case, they progress at rate $\alpha$ from one sub-state of $E_k^{(\mathrm{V})}$, $P_k^{(\mathrm{V})}$, $I_k^{(\mathrm{V})}$, $L_k^{(\mathrm{V})}$ to the corresponding counterpart reflecting no immunization, partial immunization, or ADE, or they recover ($R$). For example, from state $E_k^{(\mathrm{V})}$ individuals progress to $E_k^{(\mathrm{NI})}$, $E_k^{(\mathrm{PI})}$, $E_k^{(\mathrm{ADE})}$, or $R$, at rates, $\alpha f_E^{(\mathrm{NI})}$, $\alpha f_E^{(\mathrm{PI})}$, and $\alpha f_E^{(\mathrm{ADE})}$, and $\alpha f_E^{(R)}$, respectively, where $f_E^{(\mathrm{NI})}$, $f_E^{(\mathrm{PI})}$, $f_E^{(\mathrm{ADE})}$, and $f_E^{(R)}$ are the fractions of individuals for whom the vaccination outcome manifests during the latent phase and leads to no immunity, partial immunity, ADE, or recovery ($f_E^{(\mathrm{NI})} + f_E^{(\mathrm{PI})} + f_E^{(\mathrm{ADE})} + f_E^{(R)} = 1$). The respective fractions in the other phases of the infections are denoted by ($f_\cdot^{(\mathrm{NI})}$, $f_\cdot^{(\mathrm{PI})}$, $f_\cdot^{(\mathrm{ADE})}$, and $f_\cdot^{(R)}$) where the subscript stands for prodromal ($P$), fully infectious ($I$), and late infectious ($L$), respectively.

**Susceptibles waiting for the vaccine upon infection**

If susceptibles waiting to be vaccinated get infected, they might not be vaccinated during their COVID-19 episode or they might be vaccinated as long as their infection remains asymptomatic. In the first case, they are moved into compartment $E_1^{(\mathrm{U})}$ and further progress through the compartments $E_k^{(\mathrm{U})}$, $P_k^{(\mathrm{U})}$ (see S1 Fig, yellow compartments in the middle). From compartment $P_{n_P}^{(\mathrm{U})}$ they progress to compartment $I_1^{(\mathrm{U},+)}$ if the infection will turn symptomatic or will be detected – this is the case for a fraction $f_{\mathrm{Sick}} + (1 - f_{\mathrm{Sick}}) f_I^{(\mathrm{U},\,+)}$ (see S1 Fig, blue compartments), where $f_I^{(\mathrm{U},\,+)}$ is the fraction of asymptomatic individuals that will be tested positive before they would receive a vaccine. The remaining fraction $1 - f_{\mathrm{Sick}} - (1 - f_{\mathrm{Sick}}) f_I^{(\mathrm{U},\,+)}$ is moved into $I_1^{(\mathrm{U},-)}$. From $I_1^{(\mathrm{U},+)}$ or $I_1^{(\mathrm{U},-)}$ individuals progress through compartments $I_k^{(\mathrm{U},+)}$ and $L_k^{(\mathrm{U},+)}$ or $I_k^{(\mathrm{U},-)}$ and $L_k^{(\mathrm{U},-)}$, respectively, before they recover ($R$) or die ($D$). Asymptomatic infections lead to recovery, whereas symptomatic infections can be lethal. The fraction of individuals in $L_{n_L}^{(\mathrm{U},\,+)}$ that dies is $f_{\mathrm{Sick}}^{(\mathrm{U},+)} f_{\mathrm{Dead}}$, where

$$f_{\mathrm{Sick}}^{(\mathrm{U},+)} = \frac{f_{\mathrm{Sick}}}{f_{\mathrm{Sick}} + (1 - f_{\mathrm{Sick}}) f_I^{(\mathrm{U},\,+)}}. \tag{3}$$

In the second case, individuals are vaccinated during the infection. They progress initially through the same compartments as in the first case, but ultimately leave them. Namely, individuals in each sub-state are removed at rate $\nu$ into an equivalent sub-state in which the effect of the vaccine is pending. The exception are the compartments $I_k^{(\mathrm{U},+)}$ and $L_k^{(\mathrm{U},+)}$, which account only for individuals who will not be vaccinated because their infection is detected or symptomatic. If vaccinated during the early phases of the infection, we treat these individuals like those that were infected while immunization was pending. More precisely, in the latent phase, individuals are moved from $E_k^{(\mathrm{U})}$ to $E_k^{(\mathrm{V})}$. Likewise, in the prodromal phase they are moved from $P_k^{(\mathrm{U})}$ to $P_k^{(\mathrm{V})}$. From there the infection progresses as described above (see Vaccinated individuals that get infected while the immune response is still pending). If individuals are vaccinated during the fully infective period it is assumed that the vaccine has no (or a slightly beneficial) or a harmful effect. The effect of the vaccine manifests as before after an average duration of $D_A$ days. If vaccinated during the fully infectious phase, individuals are moved into the corresponding compartment $I_k^{(I,\mathrm{V})}$ (see S1 Fig, grey compartments on the left). As long as the vaccine shows no effect they further progress through the states $I_k^{(I,\mathrm{V})}$ and $L_k^{(I,\mathrm{V})}$ before recovery ($R$) – note these infections are asymptomatic and will not result in death. If the vaccine does not alter the course of the infection, which occurs with probability $f_I^{(I,\sim)}$ and $f_L^{(I,\sim)}$ if the outcome of the vaccination manifests during the fully infectious or late infectious phase, respectively, individuals are moved from one of the compartments $I_k^{(I,\mathrm{V})}$ or $L_k^{(I,\mathrm{V})}$ to a matching compartment $I_k^{(I,\sim)}$ or $L_k^{(I,\sim)}$, through which individuals progress similarly. However, if the vaccine has a deleterious effect, individuals are analogously moved in the sub-states $I_k^{(I,*)}$ or $L_k^{(I,*)}$. Progression through these states follows again the same pattern (see S1 Fig). However, at the end of the infection a fraction $f_{\mathrm{Sick}}^{(I,*)} f_{\mathrm{Dead}}^{(I,*)}$ dies ($D$), and the rest recovers ($R$). Asymptomatic individuals vaccinated in the late infectious phase are moved from a sub-state $L_k^{(\mathrm{U},-)}$ to sub-state $L_k^{(L,\mathrm{V})}$. From there individuals progress to the remaining states $L_k^{(L,\mathrm{V})}$ and ultimately $R$, if the effect of the vaccine is pending before recovery. If the vaccine has no (or a slightly beneficial) effect, which happens with probability $f_L^{(L,\sim)}$, individuals are moved to states $L_k^{(L,\sim)}$ through which they progress before reaching $R$. If the vaccine has a deleterious effect, individuals are moved from $L_k^{(L,\mathrm{V})}$ to $L_k^{(L,*)}$ and progress through these states before recovery ($R$) or death ($D$) – here a fraction $f_{\mathrm{Sick}}^{(L,*)} f_{\mathrm{Dead}}^{(L,*)}$ of infections is lethal.

#### Symptomatic infections

The absolute numbers of symptomatic infections in the fully infectious phase are

$$I_{\mathrm{Sick}}^{(\mathrm{NI})}(t) = f_{\mathrm{Sick}} I_{\mathrm{Sum}}^{(\mathrm{NI})}(t), \tag{4a}$$

among individuals that failed to immunize or are unvaccinable,

$$I_{\mathrm{Sick}}^{(\mathrm{PI})}(t) = f_{\mathrm{Sick}}^{(\mathrm{PI})} I_{\mathrm{Sum}}^{(\mathrm{PI})}(t), \tag{4b}$$

among individuals that partially immunized,

$$I_{\mathrm{Sick}}^{(\mathrm{ADE})}(t) = f_{\mathrm{Sick}}^{(\mathrm{ADE})} I_{\mathrm{Sum}}^{(\mathrm{ADE})}(t), \tag{4c}$$

among individuals that developed ADE,

$$I_{\mathrm{Sick}}^{(\mathrm{V})}(t) = f_{\mathrm{Sick}} I_{\mathrm{Sum}}^{(\mathrm{V})}(t), \tag{4d}$$

among individuals with pending effect of the vaccination,

$$I_{\mathrm{Sick}}^{(\mathrm{U},+)}(t) = f_{\mathrm{Sick}}^{(\mathrm{U},+)} I_{\mathrm{Sum}}^{(\mathrm{U},+)}(t), \tag{4e}$$

among individuals that waited to be vaccinated but became symptomatic or were diagnosed with COVID-19, and

$$I_{\mathrm{Sick}}^{(I,*)}(t) = f_{\mathrm{Sick}}^{(I,*)} I_{\mathrm{Sum}}^{(I,*)}(t), \tag{4f}$$

among individuals vaccinated during the fully infectious period. Similarly, the numbers of symptomatic infections in the late infectious phase are

$$L^{(\mathrm{NI})}_{\mathrm{Sick}}(t) = f_{\mathrm{Sick}} L^{(\mathrm{NI})}_{\mathrm{Sum}}(t), \tag{5a}$$

among individuals that failed to immunize or are unvaccinable,

$$L^{(\mathrm{PI})}_{\mathrm{Sick}}(t) = f^{(\mathrm{PI})}_{\mathrm{Sick}} L^{(\mathrm{PI})}_{\mathrm{Sum}}(t), \tag{5b}$$

among individuals that partially immunized,

$$L^{(\mathrm{ADE})}_{\mathrm{Sick}}(t) = f^{(\mathrm{ADE})}_{\mathrm{Sick}} L^{(\mathrm{ADE})}_{\mathrm{Sum}}(t), \tag{5c}$$

among individuals that developed ADE,

$$L^{(\mathrm{V})}_{\mathrm{Sick}}(t) = f_{\mathrm{Sick}} L^{(\mathrm{V})}_{\mathrm{Sum}}(t), \tag{5d}$$

among individuals with pending effect of the vaccination,

$$L^{(\mathrm{U},+)}_{\mathrm{Sick}}(t) = f^{(\mathrm{U},+)}_{\mathrm{Sick}} L^{(\mathrm{U},+)}_{\mathrm{Sum}}(t), \tag{5e}$$

among individuals that waited to be vaccinated but became symptomatic or were diagnosed with COVID-19,

$$L^{(I,*)}_{\mathrm{Sick}}(t) = f^{(I,*)}_{\mathrm{Sick}} L^{(I,*)}_{\mathrm{Sum}}(t), \tag{5f}$$

among individuals vaccinated during the fully infectious period, and

$$L^{(L,*)}_{\mathrm{Sick}}(t) = f^{(L,*)}_{\mathrm{Sick}} L^{(L,*)}_{\mathrm{Sum}}(t), \tag{5g}$$

among individuals vaccinated during the late infectious period.

#### Contact rate

The basic reproduction number, $R_0$ is defined as the average number of secondary infections per primary infection in a virgin population in the absence of any interventions. This number fluctuates seasonally around its annual average $\bar{R}_0$. It is modelled as

$$R_0(t) := \bar{R}_0 \left(1 + a \cos\left(2\pi \frac{t - t_{R_{0_{\max}}}}{365}\right)\right), \tag{6}$$

where, $a$ ($0 \leq a \leq 1$) is the amplitude mediating the seasonal effect on disease transmission, and $t_{R_{0_{\max}}}$ represents the time of the year when $R_0$ reaches its maximum.

Susceptible individuals get infected by random contacts with infected individuals that can transmit the disease and are not isolated. the relative contagiousness during the prodromal, fully infectious, and late infectious periods are respectively $c_P$, $c_I$, and $c_L$. Therefore, the effective contact rates in the three respective phases are

$$\boldsymbol{\beta}_P(t) := \frac{c_P R_0(t)}{c_P D_P + c_I D_I + c_L D_L}, \tag{7a}$$

$$\boldsymbol{\beta}_I(t) := \frac{c_I R_0(t)}{c_P D_P + c_I D_I + c_L D_L}, \tag{7b}$$

and

$$\boldsymbol{\beta}_L(t) := \frac{c_L R_0(t)}{c_P D_P + c_I D_I + c_L D_L}. \tag{7c}$$

The effective number of individuals that can infect susceptibles is determined by case isolation (i.e., quarantine or home isolation). The contact rates are mediated additionally by general contact reduction.

#### Case isolation

During the time interval $[t_{\text{Iso}_1}, t_{\text{Iso}_2}]$, a fraction $f_{\text{Iso}}$ of individuals with symptomatic infections that seek medical help will be isolated in quarantine wards until the wards are full. Then they are sent into home isolation. Quarantine wards guarantee perfect isolation, whereas home isolation reduces only a fraction $p_{\text{Home}}$ of contacts. Those individuals that are waiting to be vaccinated but diagnosed with COVID-19, i.e., the asymptomatic infections in the compartments $I_k^{(\text{U},+)}$ and $L_k^{(\text{U},+)}$, are also isolated. Therefore, the total number of individuals that is isolated in quarantine wards or at home at time $t$ is

$$\begin{aligned} Q(t) =& f_{\text{Iso}} \Big( I_{\text{Sick}}^{(\text{V})}(t) + L_{\text{Sick}}^{(\text{V})}(t) + I_{\text{Sick}}^{(\text{NI})}(t) + L_{\text{Sick}}^{(\text{NI})}(t) + I_{\text{Sick}}^{(\text{PI})}(t) + L_{\text{Sick}}^{(\text{PI})}(t) \\ &+ I_{\text{Sick}}^{(\text{ADE})}(t) + L_{\text{Sick}}^{(\text{ADE})}(t) + I_{\text{Sick}}^{(I,*)}(t) + L_{\text{Sick}}^{(I,*)}(t) + L_{\text{Sick}}^{(L,*)}(t) \Big) \\ &+ \Big(1 - f_{\text{Sick}}^{(\text{U},+)}\Big) \Big( I_{\text{Sum}}^{(\text{U},+)}(t) + L_{\text{Sum}}^{(\text{U},+)}(t) \Big). \end{aligned} \tag{8}$$

The total numbers of fully infectious individuals, whose immunization is pending (V), that are unvaccinable or failed to immunized (NI), were partially immunized (PI), developed ADE, or were vaccinated while being fully infectious $(I, *)$, in quarantine are

$$I_{\text{Iso}}^{(\cdot)} = \begin{cases} f_{\text{Iso}} I_{\text{Sick}}^{(\cdot)} & \text{if } t_{\text{Iso}_1} \le t \le t_{\text{Iso}_2} \text{ and } Q(t) \le Q_{\max}, \\ f_{\text{Iso}} I_{\text{Sick}}^{(\cdot)} \dfrac{Q_{\max}}{Q(t)} & \text{if } t_{\text{Iso}_1} \le t \le t_{\text{Iso}_2} \text{ and } Q(t) > Q_{\max}, \\ 0 & \text{otherwise}, \end{cases} \tag{9a}$$

where "." is a placeholder for "V", "NI", "PI", "ADE", and "$I, *$". Furthermore, the number of fully infectious individuals that waited to be vaccinated but became symptomatic or were diagnosed with COVID-19 in isolation is

$$I_{\text{Iso}}^{(\text{U},+)} = \begin{cases} \left(f_{\text{Sick}}^{(\text{U},+)} f_{\text{Iso}} + \left(1 - f_{\text{Sick}}^{(\text{U},+)}\right)\right) I_{\text{Sum}}^{(\text{U},+)}(t) & \\ & \text{if } t_{\text{Iso}_1} \leq t \leq t_{\text{Iso}_2} \text{ and } Q(t) \leq Q_{\max}, \\ \left(f_{\text{Sick}}^{(\text{U},+)} f_{\text{Iso}} + \left(1 - f_{\text{Sick}}^{(\text{U},+)}\right)\right) I_{\text{Sum}}^{(\text{U},+)}(t) \dfrac{Q_{\max}}{Q(t)} & \\ & \text{if } t_{\text{Iso}_1} \leq t \leq t_{\text{Iso}_2} \text{ and } Q(t) > Q_{\max}, \\ 0 & \text{otherwise.} \end{cases} \tag{9b}$$

The numbers of fully infectious individuals in home isolation in the different groups are

$$I_{\text{Home}}^{(.)} = \begin{cases} I_{\text{Sick}}^{(.)} f_{\text{Iso}} \left(1 - \dfrac{Q_{\max}}{Q(t)}\right) & \text{if } t_{\text{Iso}_1} \leq t \leq t_{\text{Iso}_2} \text{ and } Q(t) > Q_{\max}, \\ 0 & \text{otherwise,} \end{cases} \tag{10a}$$

where "." is a placeholder for "V", "NI", "PI", "ADE", and "$I, *$". Furthermore, the number of fully infectious individuals that waited to be vaccinated but became symptomatic or were diagnosed with COVID-19 in home isolation is

$$I_{\text{Home}}^{(\text{U},+)}(t) = \begin{cases} \left(f_{\text{Sick}}^{(\text{U},+)} f_{\text{Iso}} + \left(1 - f_{\text{Sick}}^{(\text{U},+)}\right)\right) I_{\text{Sum}}^{(\text{U},+)}(t) \left(1 - \dfrac{Q_{\max}}{Q(t)}\right) & \\ & \text{if } t_{\text{Iso}_1} \leq t \leq t_{\text{Iso}_2} \text{ and } Q(t) > Q_{\max}, \\ 0 & \text{otherwise.} \end{cases} \tag{10b}$$

Therefore, the effective numbers of fully infectious individuals in those groups subject to isolation that can infect susceptibles becomes

$$I_{\text{Eff}}^{(.)} = I_{\text{Sum}}^{(.)} - I_{\text{Iso}}^{(.)} - (1 - p_{\text{Home}}) I_{\text{Home}}^{(.)}, \tag{11}$$

where "." is a placeholder for "V", "NI", "PI", "ADE", "$I, *$", "U,+".

The total numbers of late infectious individuals, whose immunization is pending (V), that are unvaccinable or failed to immunized (NI), were partially immunized (PI), developed ADE, or were vaccinated while being fully or late infectious ($I, *$, or $L, *$), in quarantine are

$$L_{\text{Iso}}^{(.)}(t) = \begin{cases} f_{\text{Iso}} L_{\text{Sick}}^{(.)} & \text{if } \quad t_{\text{Iso}_1} \leq t \leq t_{\text{Iso}_2} \text{ and } Q(t) \leq Q_{\max}, \\ f_{\text{Iso}} L_{\text{Sick}}^{(.)} \dfrac{Q_{\max}}{Q(t)} & \text{if } t_{\text{Iso}_1} \leq t \leq t_{\text{Iso}_2} \text{ and } Q(t) > Q_{\max}, \\ 0 & \text{otherwise,} \end{cases} \tag{12a}$$

where "." is a placeholder for "V", "NI", "PI", "ADE", "$I, *$" and "$L, *$". Furthermore, the number of late infectious individuals that waited to be vaccinated but became symptomatic or were diagnosed with COVID-19 in isolation is

$$L_{\mathrm{Iso}}^{(\mathrm{U},+)}(t) = \begin{cases} \left(f_{\mathrm{Sick}}^{(\mathrm{U},+)} f_{\mathrm{Iso}} + \left(1 - f_{\mathrm{Sick}}^{(\mathrm{U},+)}\right)\right) L_{\mathrm{Sum}}^{(\mathrm{U},+)}(t) & \\ \qquad \text{if } t_{\mathrm{Iso}_1} \leq t \leq t_{\mathrm{Iso}_2} \text{ and } Q(t) \leq Q_{\max}, & \\ \left(f_{\mathrm{Sick}}^{(\mathrm{U},+)} f_{\mathrm{Iso}} + \left(1 - f_{\mathrm{Sick}}^{(\mathrm{U},+)}\right)\right) L_{\mathrm{Sum}}^{(\mathrm{U},+)}(t) \dfrac{Q_{\max}}{Q(t)} & \\ \qquad \text{if } t_{\mathrm{Iso}_1} \leq t \leq t_{\mathrm{Iso}_2} \text{ and } Q(t) > Q_{\max}, & \\ 0 & \text{otherwise.} \end{cases} \tag{12b}$$

The numbers of late infectious individuals in home isolation in different groups are

$$L_{\mathrm{Home}}^{(.)}(t) = \begin{cases} L_{\mathrm{Sick}}^{(.)} f_{\mathrm{Iso}} \left(1 - \dfrac{Q_{\max}}{Q(t)}\right) & \text{if } t_{\mathrm{Iso}_1} \leq t \leq t_{\mathrm{Iso}_2} \text{ and } Q(t) > Q_{\max}, \\ 0 & \text{otherwise,} \end{cases} \tag{13a}$$

where "." is a placeholder for "V", "NI", "PI", "ADE", "$I,*$", and "$L,*$". Furthermore, the number of late infectious individuals that waited to be vaccinated but became symptomatic or were diagnosed with COVID-19 in home isolation is

$$L_{\mathrm{Home}}^{(\mathrm{U},+)}(t) = \begin{cases} \left(f_{\mathrm{Sick}}^{(\mathrm{U},+)} f_{\mathrm{Iso}} + \left(1 - f_{\mathrm{Sick}}^{(\mathrm{U},+)}\right)\right) L_{\mathrm{Sum}}^{(\mathrm{U},+)}(t) \left(1 - \dfrac{Q_{\max}}{Q(t)}\right) & \\ \qquad \text{if } t_{\mathrm{Iso}_1} \leq t \leq t_{\mathrm{Iso}_2} \text{ and } Q(t) > Q_{\max}, & \\ 0 & \text{otherwise.} \end{cases} \tag{13b}$$

The effective numbers of late infectious individuals in those groups subject to isolation that can infect susceptibles becomes

$$L_{\mathrm{Eff}}^{(.)}(t) = L_{\mathrm{Sum}}^{(.)} - L_{\mathrm{Iso}}^{(.)}(t) - (1 - p_{\mathrm{Home}}) L_{\mathrm{Home}}^{(.)}(t), \tag{14}$$

where "." is a placeholder for "V", "NI", "PI", "ADE", "$I,*$", "U,+".

#### General contact reduction

During designated time intervals general contact reduction (curfews, social distancing, cancellation of mass events, etc.) is sustained. Hence the number of contacts at time $t$ is reduced by a fraction $p_{\mathrm{Dist}}$. Here, we assume an initial phase without general contact reduction until time $t_{\mathrm{Dist}_1}$, a "hard lockdown" between $t_{\mathrm{Dist}_1}$ and $t_{\mathrm{Dist}_2}$, followed by a phase of "relief" between $t_{\mathrm{Dist}_2}$ and $t_{\mathrm{Dist}_3}$, a "soft lockdown" between $t_{\mathrm{Dist}_3}$ and $t_{\mathrm{Dist}_4}$, a second "hard lockdown" between $t_{\mathrm{Dist}_4}$ and $t_{\mathrm{Dist}_5}$, and finally a second phase of "relief" until $t_{\text{Dist}_5}$, after which general contact reduction is lifted, resulting in

$$p_{\text{Dist}} = \begin{cases} p_{\text{Cont}_1} & \text{for } t_{\text{Dist}_1} \le t \le t_{\text{Dist}_2}, \\ p_{\text{Cont}_2} & \text{for } t_{\text{Dist}_2} \le t \le t_{\text{Dist}_3}, \\ p_{\text{Cont}_3} & \text{for } t_{\text{Dist}_3} \le t \le t_{\text{Dist}_4}, \\ p_{\text{Cont}_4} & \text{for } t_{\text{Dist}_4} \le t \le t_{\text{Dist}_5}, \\ p_{\text{Cont}_5} & \text{for } t_{\text{Dist}_5} \le t \le t_{\text{Dist}_6}, \\ 0 & \text{otherwise.} \end{cases} \tag{15}$$

Note, that the function $p_{\text{Dist}}$ can be adjusted as necessary.

### Force of infection

Taking the contact rates (7), general contact reduction (15), and the numbers of infectious individuals that effectively participate in infection (11), (14) into account, the force of infection becomes

$$\begin{aligned} \lambda(t) = \big(1 - p_{\text{Cont}}(t)\big)\Big(&\beta_P(t)\Big(P^{(\text{U})}_{\text{Sum}}(t) + P^{(\text{V})}_{\text{Sum}}(t) + P^{(\text{NI})}_{\text{Sum}}(t) + P^{(\text{PI})}_{\text{Sum}}(t) + P^{(\text{ADE})}_{\text{Sum}}(t)\Big) \\ &+\beta_I(t)\Big(I^{(\text{U},-)}_{\text{Sum}}(t) + I^{(\text{U},+)}_{\text{Eff}}(t) + I^{(\text{V})}_{\text{Eff}}(t) + I^{(\text{NI})}_{\text{Eff}}(t) + I^{(\text{PI})}_{\text{Eff}}(t) \\ &\quad + I^{(\text{ADE})}_{\text{Eff}}(t) + I^{(I,\text{V})}_{\text{Sum}}(t) + I^{(I,\sim)}_{\text{Sum}}(t) + I^{(I,*)}_{\text{Eff}}(t)\Big) \\ &+\beta_L(t)\Big(L^{(\text{U},-)}_{\text{Sum}}(t) + L^{(\text{U},+)}_{\text{Eff}}(t) + L^{(\text{V})}_{\text{Eff}}(t) + L^{(\text{NI})}_{\text{Eff}}(t) + L^{(\text{PI})}_{\text{Eff}}(t) \\ &\quad + L^{(\text{ADE})}_{\text{Eff}}(t) + L^{(I,\text{V})}_{\text{Sum}}(t) + L^{(I,\sim)}_{\text{Sum}}(t) + L^{(I,*)}_{\text{Eff}}(t) \\ &\quad + L^{(L,\text{V})}_{\text{Sum}}(t) + L^{(L,\sim)}_{\text{Sum}}(t) + L^{(L,*)}_{\text{Eff}}(t)\Big)\Big) + \lambda_{\text{Ext}}, \end{aligned} \tag{16}$$

where $\lambda_{\text{Ext}}$ is the external force of infection, i.e., caused by infections from outside the population. Individuals that developed partial immunity or ADE are protected from infection by some amount. Namely, the infectiousness at a given contact is reduced by a fractions $p_{\text{PI}}$ and $p_{\text{ADE}}$, respectively. Thus, the forces of infection experience by susceptible individuals with partial immunity and ADE are, respectively, $p_{\text{PI}}\lambda(t)$ and $p_{\text{ADE}}\lambda(t)$.

### Model dynamics

The model is formulated as a system of differential equations following the flow chart in S1 Fig.

#### Dynamics of susceptibles

The change in the number of unvaccinable susceptibles is determined only by the force of infection and given by

$$\frac{dS^{(\text{NV})}(t)}{dt} = -\lambda(t)\frac{S^{(\text{NV})}(t)}{N}. \tag{17a}$$

The number of susceptibles in line for the vaccine are reduced by those that get infected or vaccinated (at rate $\nu$). Hence, their dynamics change according to

$$\frac{dS^{(\mathrm{U})}(t)}{dt} = -\lambda(t)\frac{S^{(\mathrm{U})}(t)}{N} - \nu S^{(\mathrm{U})}(t). \tag{17b}$$

Once susceptibles are vaccinated, their vaccination outcome is pending for an average duration of $D_A = 1/\alpha$. During this time they can be infected. Hence, their number changes according to

$$\frac{dS^{(\mathrm{V})}(t)}{dt} = \nu S^{(\mathrm{U})}(t) - \lambda(t)\frac{S^{(\mathrm{V})}(t)}{N} - \alpha S^{(\mathrm{V})}(t). \tag{17c}$$

After the average duration $D_A$, fractions $f_S^{(\mathrm{NI})}$, $f_S^{(\mathrm{PI})}$, and $f_S^{(\mathrm{ADE})}$ of susceptibles fail to immunize, develop partial immunity or ADE, respectively. Those individuals that failed to immunize are equally susceptible as unvaccinable individuals, whereas the susceptibility of those with partial immunity or ADE reduce by the fractions $p_{\mathrm{PI}}\lambda(t)$ and $p_{\mathrm{ADE}}\lambda(t)$. The numbers in these three categories of susceptibles hence changes according to

$$\frac{dS^{(\mathrm{NI})}(t)}{dt} = \alpha f_S^{(\mathrm{NI})} S^{(\mathrm{V})}(t) - \lambda(t)\frac{S^{(\mathrm{NI})}(t)}{N}, \tag{17d}$$

$$\frac{dS^{(\mathrm{PI})}(t)}{dt} = \alpha f_S^{(\mathrm{PI})} S^{(\mathrm{V})}(t) - p_{\mathrm{PI}}\lambda(t)\frac{S^{(\mathrm{PI})}(t)}{N}, \tag{17e}$$

and

$$\frac{dS^{(\mathrm{ADE})}(t)}{dt} = \alpha f_S^{(\mathrm{ADE})} S^{(\mathrm{V})}(t) - p_{\mathrm{ADE}}\lambda(t)\frac{S^{(\mathrm{ADE})}(t)}{N}. \tag{17f}$$

##### Dynamics of latent-infected individuals

Upon infection susceptibles become latently infected. Latent-infected individuals progress through sub-states at rate $\varepsilon$. As described above, infections of unvaccinable individuals and those that failed to immunize are indistinguishable and are modelled with the same compartments. The dynamics of latent-infected unvaccinable individuals and those that failed to immunize are

$$\frac{dE_1^{(\mathrm{NI})}(t)}{dt} = \lambda(t)\frac{S^{(\mathrm{NV})}(t) + S^{(\mathrm{NI})}(t)}{N} + \alpha f_E^{(\mathrm{NI})} E_1^{(\mathrm{V})}(t) - \varepsilon E_1^{(\mathrm{NI})}(t), \tag{18a}$$

$$\frac{dE_k^{(\mathrm{NI})}(t)}{dt} = \varepsilon E_{k-1}^{(\mathrm{NI})}(t) + \alpha f_E^{(\mathrm{NI})} E_k^{(\mathrm{V})}(t) - \varepsilon E_k^{(\mathrm{NI})}(t) \qquad \text{for } 2 \leq k \leq n_E. \tag{18b}$$

Those latent-infected individuals in line for the vaccine, still get vaccinated at rate $\nu$, hence their dynamics change according to

$$\frac{dE_1^{(\mathrm{U})}(t)}{dt} = \lambda(t)\frac{S^{(\mathrm{U})}(t)}{N} - \nu E_1^{(\mathrm{U})}(t) - \varepsilon E_1^{(\mathrm{U})}(t), \tag{18c}$$

$$\frac{dE_k^{(\mathrm{U})}(t)}{dt} = \varepsilon E_{k-1}^{(\mathrm{U})}(t) - \varepsilon E_k^{(\mathrm{U})}(t) - \nu E_k^{(\mathrm{U})}(t) \qquad \text{for } 2 \leq k \leq n_E, \tag{18d}$$

The number of latent-infected individuals with pending vaccination outcome increase by those that get vaccinated during the latent phase and decrease by those for whom the outcome manifests. Hence their dynamics change according to

$$\frac{dE_1^{(\mathrm{V})}(t)}{dt} = \lambda(t)\frac{S^{(\mathrm{V})}(t)}{N} + \nu E_1^{(\mathrm{U})}(t) - \varepsilon E_1^{(\mathrm{V})}(t) - \alpha E_1^{(\mathrm{V})}(t), \tag{18e}$$

$$\frac{dE_k^{(\mathrm{V})}(t)}{dt} = \varepsilon E_{k-1}^{(\mathrm{V})}(t) + \nu E_k^{(\mathrm{U})}(t) - \varepsilon E_k^{(\mathrm{V})}(t) - \alpha E_k^{(\mathrm{V})}(t) \qquad \text{for } 2 \le k \le n_E. \tag{18f}$$

The vaccination outcome can be a failure to immunize (cf. Eq. (18a), (18b)), a partial immunity, or ADE. The number of latent-infected individuals with respectively partial immunity, or ADE changes according to

$$\frac{dE_1^{(\mathrm{PI})}(t)}{dt} = p_{\mathrm{PI}}\lambda(t)\lambda(t)\frac{S^{(\mathrm{PI})}(t)}{N} + \alpha f_E^{(\mathrm{PI})} E_1^{(\mathrm{V})}(t) - \varepsilon E_1^{(\mathrm{PI})}(t), \tag{18g}$$

$$\frac{dE_k^{(\mathrm{PI})}(t)}{dt} = \varepsilon E_{k-1}^{(\mathrm{PI})}(t) + \alpha f_E^{(\mathrm{PI})} E_k^{(\mathrm{V})}(t) - \varepsilon E_k^{(\mathrm{PI})}(t) \qquad \text{for } 2 \le k \le n_E, \tag{18h}$$

$$\frac{dE_1^{(\mathrm{ADE})}(t)}{dt} = p_{\mathrm{ADE}}\lambda(t)\frac{S^{(\mathrm{ADE})}(t)}{N} + \alpha f_E^{(\mathrm{ADE})} E_1^{(\mathrm{V})}(t) - \varepsilon E_1^{(\mathrm{ADE})}(t), \tag{18i}$$

and

$$\frac{dE_k^{(\mathrm{ADE})}(t)}{dt} = \varepsilon E_{k-1}^{(\mathrm{ADE})}(t) + \alpha f_E^{(\mathrm{ADE})} E_k^{(\mathrm{V})}(t) - \varepsilon E_k^{(\mathrm{ADE})}(t) \qquad \text{for } 2 \le k \le n_E. \tag{18j}$$

**Dynamics of prodromal infections**

The dynamics of the prodromal-infected individuals are analogous to those of the latent-infected individuals, but they progress through sub-states at rate $\varphi$. The dynamics of prodromal individuals in the various groups hence become

$$\frac{dP_1^{(\mathrm{U})}(t)}{dt} = \varepsilon E_{n_E}^{(\mathrm{U})}(t) - \varphi P_1^{(\mathrm{U})}(t) - \nu P_1^{(\mathrm{U})}(t), \tag{19a}$$

$$\frac{dP_k^{(\mathrm{U})}(t)}{dt} = \varphi P_{k-1}^{(\mathrm{U})}(t) - \varphi P_k^{(\mathrm{U})}(t) - \nu P_k^{(\mathrm{U})}(t) \qquad \text{for } 2 \le k \le n_P, \tag{19b}$$

$$\frac{dP_1^{(\mathrm{V})}(t)}{dt} = \varepsilon E_{n_E}^{(\mathrm{V})}(t) + \nu P_1^{(\mathrm{U})}(t) - \varphi P_1^{(\mathrm{V})}(t) - \alpha P_1^{(\mathrm{V})}(t), \tag{19c}$$

$$\frac{dP_k^{(\mathrm{V})}(t)}{dt} = \varphi P_{k-1}^{(\mathrm{V})}(t) + \nu P_k^{(\mathrm{U})}(t) - \varphi P_k^{(\mathrm{V})}(t) - \alpha P_k^{(\mathrm{V})}(t) \qquad \text{for } 2 \le k \le n_P, \tag{19d}$$

$$\frac{dP_1^{(\mathrm{NI})}(t)}{dt} = \varepsilon E_{n_E}^{(\mathrm{NI})}(t) + \alpha f_P^{(\mathrm{NI})} P_1^{(\mathrm{V})}(t) - \varphi P_1^{(\mathrm{NI})}(t), \tag{19e}$$

$$\frac{dP_k^{(\mathrm{NI})}(t)}{dt} = \varphi P_{k-1}^{(\mathrm{NI})}(t) + \alpha f_P^{(\mathrm{NI})} P_k^{(\mathrm{V})}(t) - \varphi P_k^{(\mathrm{NI})}(t) \qquad \text{for } 2 \le k \le n_P, \tag{19f}$$

$$\frac{dP_1^{(\mathrm{PI})}(t)}{dt} = \varepsilon E_{n_E}^{(\mathrm{PI})}(t) + \alpha f_P^{(\mathrm{PI})} P_1^{(\mathrm{V})}(t) - \varphi P_1^{(\mathrm{PI})}(t), \tag{19g}$$

$$\frac{dP_k^{(\mathrm{PI})}(t)}{dt} = \varphi P_{k-1}^{(\mathrm{PI})}(t) + \alpha f_P^{(\mathrm{PI})} P_k^{(\mathrm{V})}(t) - \varphi P_k^{(\mathrm{PI})}(t) \qquad \text{for } 2 \le k \le n_P, \tag{19h}$$

$$\frac{dP_1^{(\mathrm{ADE})}(t)}{dt} = \varepsilon E_{n_E}^{(\mathrm{ADE})}(t) + \alpha f_P^{(\mathrm{ADE})} P_1^{(\mathrm{V})}(t) - \varphi P_1^{(\mathrm{ADE})}(t), \tag{19i}$$

and

$$\frac{dP_k^{(\mathrm{ADE})}(t)}{dt} = \varphi P_{k-1}^{(\mathrm{ADE})}(t) + \alpha f_P^{(\mathrm{ADE})} P_k^{(\mathrm{V})}(t) - \varphi P_k^{(\mathrm{ADE})}(t) I_k^{(\mathrm{V})}(t) \text{ for } 2 \leq k \leq n_P. \quad (19j)$$

##### Dynamics of fully infectious individuals

The dynamics of fully infectious individuals that failed to immunize or are unvaccinable, developed partial immunity, or ADE are analogous to their respective dynamics in the prodromal phase, only that they progress through sub-states at a rate $\gamma$. The same is true for vaccinated fully infectious individuals, whose vaccination outcome is pending, however, fully infectious individuals that are being vaccinated are not moved to their compartments, but to separate ones. Hence, the dynamics of the mentioned groups of fully infectious individuals become

$$\frac{dI_1^{(\mathrm{V})}(t)}{dt} = \varphi P_{n_P}^{(\mathrm{V})}(t) - \gamma I_1^{(\mathrm{V})}(t) - \alpha I_1^{(\mathrm{V})}(t), \quad (20a)$$

$$\frac{dI_k^{(\mathrm{V})}(t)}{dt} = \gamma I_{k-1}^{(\mathrm{V})}(t) - \gamma I_k^{(\mathrm{V})}(t) - \alpha I_k^{(\mathrm{V})}(t) \quad \text{for } 2 \leq k \leq n_I, \quad (20b)$$

$$\frac{dI_1^{(\mathrm{NI})}(t)}{dt} = \varphi P_{n_P}^{(\mathrm{NI})}(t) + \alpha f_I^{(\mathrm{NI})} I_1^{(\mathrm{V})}(t) - \gamma I_1^{(\mathrm{NI})}(t), \quad (20c)$$

$$\frac{dI_k^{(\mathrm{NI})}(t)}{dt} = \gamma I_{k-1}^{(\mathrm{NI})}(t) + \alpha f_I^{(\mathrm{NI})} I_k^{(\mathrm{V})}(t) - \gamma I_k^{(\mathrm{NI})}(t) \quad \text{for } 2 \leq k \leq n_I, \quad (20d)$$

$$\frac{dI_1^{(\mathrm{PI})}(t)}{dt} = \varphi P_{n_P}^{(\mathrm{PI})}(t) + \alpha f_I^{(\mathrm{PI})} I_1^{(\mathrm{V})}(t) - \gamma I_1^{(\mathrm{PI})}(t), \quad (20e)$$

$$\frac{dI_k^{(\mathrm{PI})}(t)}{dt} = \gamma I_{k-1}^{(\mathrm{PI})}(t) + \alpha f_I^{(\mathrm{PI})} I_k^{(\mathrm{V})}(t) - \gamma I_k^{(\mathrm{PI})}(t) \quad \text{for } 2 \leq k \leq n_I, \quad (20f)$$

$$\frac{dI_1^{(\mathrm{ADE})}(t)}{dt} = \varphi P_{n_P}^{(\mathrm{ADE})}(t) + \alpha f_I^{(\mathrm{ADE})} I_1^{(\mathrm{V})}(t) - \gamma I_1^{(\mathrm{ADE})}(t), \quad (20g)$$

and

$$\frac{dI_k^{(\mathrm{ADE})}(t)}{dt} = \gamma I_{k-1}^{(\mathrm{ADE})}(t) + \alpha f_I^{(\mathrm{ADE})} I_k^{(\mathrm{V})}(t) - \gamma I_k^{(\mathrm{ADE})}(t) \quad \text{for } 2 \leq k \leq n_I. \quad (20h)$$

For the unvaccinated individuals in the fully infectious phase, it is distinguished between those that still can get vaccinated ($I_k^{(\mathrm{U},-)}$ – those that remain asymptomatic and undiagnosed), and those that are not vaccinated because they are symptomatic or diagnosed ($I_k^{(\mathrm{U},+)}$). Their dynamics become

$$\frac{dI_1^{(\mathrm{U},+)}(t)}{dt} = \varphi\Big(f_{\mathrm{Sick}} + \big(1 - f_{\mathrm{Sick}}\big) f_I^{(\mathrm{U},\,+)}\Big) P_{n_P}^{(\mathrm{U})}(t) - \gamma I_1^{(\mathrm{U},+)}(t), \quad (20i)$$

$$\frac{dI_k^{(\mathrm{U},+)}(t)}{dt} = \gamma I_{k-1}^{(\mathrm{U},+)}(t) - \gamma I_k^{(\mathrm{U},+)}(t) \quad \text{for } 2 \leq k \leq n_I, \quad (20j)$$

$$\frac{dI_1^{(\mathrm{U},-)}(t)}{dt} = \varphi(1 - f_{\mathrm{Sick}})\Big(1 - f_I^{(\mathrm{U},\,+)}\Big) P_{n_P}^{(\mathrm{U})}(t) - \gamma I_1^{(\mathrm{U},-)}(t) - \nu I_1^{(\mathrm{U},-)}(t), \quad (20k)$$

and

$$\frac{dI_k^{(\mathrm{U},-)}(t)}{dt} = \gamma I_{k-1}^{(\mathrm{U},-)}(t) - \gamma I_k^{(\mathrm{U},-)}(t) - \nu I_k^{(\mathrm{U},-)}(t) \quad \text{for } 2 \leq k \leq n_I. \quad (20l)$$

The dynamics of fully infectious individuals that got vaccinated during this phase, for whom the vaccination outcome is still pending, are

$$\frac{dI_1^{(I,\mathrm{V})}(t)}{dt} = \nu I_1^{(\mathrm{U},-)}(t) - \gamma I_1^{(I,\mathrm{V})}(t) - \alpha I_1^{(I,\mathrm{V})}(t), \tag{20m}$$

and

$$\frac{dI_k^{(I,\mathrm{V})}(t)}{dt} = \gamma I_{k-1}^{(I,\mathrm{V})}(t) + \nu I_k^{(\mathrm{U},-)}(t) - \gamma I_k^{(I,\mathrm{V})}(t) - \alpha I_k^{(I,\mathrm{V})}(t) \quad \text{for } 2 \leq k \leq n_I, \tag{20n}$$

The numbers of asymptomatic fully infectious individuals that got vaccinated during this phase, but the immunization had neutral effect, changes according to

$$\frac{dI_1^{(I,\sim)}(t)}{dt} = \alpha f_I^{(I,\sim)} I_1^{(I,\mathrm{V})}(t) - \gamma I_1^{(I,\sim)}(t), \tag{20o}$$

and

$$\frac{dI_k^{(I,\sim)}(t)}{dt} = \gamma I_{k-1}^{(I,\sim)}(t) + \alpha f_I^{(I,\sim)} I_k^{(I,\mathrm{V})}(t) - \gamma I_k^{(I,\sim)}(t) \quad \text{for } 2 \leq k \leq n_I. \tag{20p}$$

The numbers of asymptomatic fully infectious individuals, vaccinated during this phase with a deleterious outcome, change according to

$$\frac{dI_1^{(I,*)}(t)}{dt} = \alpha\left(1 - f_I^{(I,\sim)}\right) I_1^{(I,\mathrm{V})}(t) - \gamma I_1^{(I,*)}(t), \tag{20q}$$

and

$$\frac{dI_k^{(I,*)}(t)}{dt} = \gamma I_{k-1}^{(I,*)}(t) + \alpha\left(1 - f_I^{(I,\sim)}\right) I_k^{(I,\mathrm{V})}(t) - \gamma I_k^{(I,*)}(t) \quad \text{for } 2 \leq k \leq n_I. \tag{20r}$$

**Dynamics of late infectious individuals**

The dynamics of late infectious individuals are analogous to those of the fully infectious. However, there are additional compartments modelling asymptomatic late infectious individuals that get vaccinated during this phase ($L_k^{(\mathrm{U},-)}$). First, the outcome of the vaccine is pending ($L_k^{(L,\mathrm{V})}$). The vaccine might ultimately have neutral effect ($L_k^{(L,\sim)}$) or a deleterious effect ($L_k^{(L,*)}$). Therefore, the numbers of late infectious individual for which the outcome of the vaccine is pending change according to

$$\frac{dL_1^{(\mathrm{V})}(t)}{dt} = \gamma I_{n_I}^{(\mathrm{V})}(t) - \delta L_1^{(\mathrm{V})}(t) - \alpha L_1^{(\mathrm{V})}(t), \tag{21a}$$

and

$$\frac{dL_k^{(\mathrm{V})}(t)}{dt} = \delta L_{k-1}^{(\mathrm{V})}(t) - \delta L_k^{(\mathrm{V})}(t) - \alpha L_k^{(\mathrm{V})}(t) \quad \text{for } 2 \leq k \leq n_L. \tag{21b}$$

Analogously to (20c)–(20h), the numbers of late infectious individuals that are unvaccinable or failed to immunize, developed partial resistance, or ADE change according to

$$\frac{dL_1^{(\mathrm{NI})}(t)}{dt} = \gamma I_{n_I}^{(\mathrm{NI})}(t) + \alpha f_L^{(\mathrm{NI})} L_1^{(\mathrm{V})}(t) - \delta L_1^{(\mathrm{NI})}(t), \tag{21c}$$

$$\frac{dL_k^{(\mathrm{NI})}(t)}{dt} = \delta L_{k-1}^{(\mathrm{NI})}(t) + \alpha f_L^{(\mathrm{NI})} L_k^{(\mathrm{V})}(t) - \delta L_k^{(\mathrm{NI})}(t), \qquad \text{for } 2 \leq k \leq n_L, \tag{21d}$$

$$\frac{dL_1^{(\mathrm{PI})}(t)}{dt} = \gamma I_{n_I}^{(\mathrm{PI})}(t) + \alpha f_L^{(\mathrm{PI})} L_1^{(\mathrm{V})}(t) - \delta L_1^{(\mathrm{PI})}(t), \tag{21e}$$

$$\frac{dL_k^{(\mathrm{PI})}(t)}{dt} = \delta L_{k-1}^{(\mathrm{PI})}(t) + \alpha f_L^{(\mathrm{PI})} L_k^{(\mathrm{V})}(t) - \delta L_k^{(\mathrm{PI})}(t) \qquad \text{for } 2 \leq k \leq n_L, \tag{21f}$$

$$\frac{dL_1^{(\mathrm{ADE})}(t)}{dt} = \gamma I_{n_I}^{(\mathrm{ADE})}(t) + \alpha f_L^{(\mathrm{ADE})} L_1^{(\mathrm{V})}(t) - \delta L_1^{(\mathrm{ADE})}(t), \tag{21g}$$

and

$$\frac{dL_k^{(\mathrm{ADE})}(t)}{dt} = \delta L_{k-1}^{(\mathrm{ADE})}(t) + \alpha f_L^{(\mathrm{ADE})} L_k^{(\mathrm{V})}(t) - \delta L_k^{(\mathrm{ADE})}(t) \qquad \text{for } 2 \leq k \leq n_L. \tag{21h}$$

The late infectious individuals that are in line for vaccination but did not get vaccinated change according to

$$\frac{dL_1^{(\mathrm{U},+)}(t)}{dt} = \gamma I_{n_I}^{(\mathrm{U},\,+)}(t) - \delta L_1^{(\mathrm{U},+)}(t), \tag{21i}$$

$$\frac{dL_k^{(\mathrm{U},+)}(t)}{dt} = \delta L_{k-1}^{(\mathrm{U},+)}(t) - \delta L_k^{(\mathrm{U},+)}(t) \qquad \text{for } 2 \leq k \leq n_L, \tag{21j}$$

$$\frac{dL_1^{(\mathrm{U},-)}(t)}{dt} = \gamma I_{n_I}^{(\mathrm{U},\,-)}(t) - \delta L_1^{(\mathrm{U},-)}(t) - \nu L_1^{(\mathrm{U},-)}(t), \tag{21k}$$

and

$$\frac{dL_k^{(\mathrm{U},-)}(t)}{dt} = \delta L_{k-1}^{(\mathrm{U},-)}(t) - \delta L_k^{(\mathrm{U},-)}(t) - \nu L_k^{(\mathrm{U},-)}(t) \qquad \text{for } 2 \leq k \leq n_L. \tag{21l}$$

Those late infectious individuals that got vaccinated during the fully infectious phase change according to

$$\frac{dL_1^{(I,\mathrm{V})}(t)}{dt} = \gamma I_{n_I}^{(I,\mathrm{V})}(t) - \delta L_1^{(I,\mathrm{V})}(t) - \alpha L_1^{(I,\mathrm{V})}(t), \tag{21m}$$

$$\frac{dL_k^{(I,\mathrm{V})}(t)}{dt} = \delta L_{k-1}^{(I,\mathrm{V})}(t) - \delta L_k^{(I,\mathrm{V})}(t) - \alpha L_k^{(I,\mathrm{V})}(t) \qquad \text{for } 2 \leq k \leq n_L, \tag{21n}$$

$$\frac{dL_1^{(I,\sim)}(t)}{dt} = \gamma I_{n_I}^{(I,\sim)}(t) + \alpha f_L^{(I,\sim)} L_1^{(I,\mathrm{V})}(t) - \delta L_1^{(I,\sim)}(t), \tag{21o}$$

$$\frac{dL_k^{(I,\sim)}(t)}{dt} = \delta L_{k-1}^{(L,\sim)}(t) + \alpha f_L^{(I,\sim)} L_k^{(I,\mathrm{V})}(t) - \delta L_k^{(I,\sim)}(t) \qquad \text{for } 2 \leq k \leq n_L, \tag{21p}$$

$$\frac{dL_1^{(I,*)}(t)}{dt} = \gamma I_{n_I}^{(I,*)}(t) + \alpha\Big(1 - f_L^{(I,\sim)}\Big) L_1^{(I,\mathrm{V})}(t) - \delta L_1^{(I,*)}(t), \tag{21q}$$

and

$$\frac{dL_k^{(I,*)}(t)}{dt} = \delta L_{k-1}^{(I,*)}(t) + \alpha\Big(1 - f_L^{(I,\sim)}\Big)L_k^{(I,\mathrm{V})}(t) - \delta L_k^{(I,*)}(t) \quad \text{for } 2 \le k \le n_L. \tag{21r}$$

Additionally, the dynamics of late infectious individuals that got vaccinated during this phase are

$$\frac{dL_1^{(L,\mathrm{V})}(t)}{dt} = \nu L_1^{(\mathrm{U},-)}(t) - \delta L_1^{(L,\mathrm{V})}(t) - \alpha L_1^{(L,\mathrm{V})}(t), \tag{21s}$$

$$\frac{dL_k^{(L,\mathrm{V})}(t)}{dt} = \delta L_{k-1}^{(L,\mathrm{V})}(t) + \nu L_k^{(\mathrm{U},-)}(t) - \delta L_k^{(L,\mathrm{V})}(t) - \alpha L_k^{(L,\mathrm{V})}(t) \quad \text{for } 2 \le k \le n_L, \tag{21t}$$

$$\frac{dL_1^{(L,\sim)}(t)}{dt} = \alpha f_L^{(L,\sim)} L_1^{(L,\mathrm{V})}(t) - \delta L_1^{(L,\sim)}(t), \tag{21u}$$

$$\frac{dL_k^{(L,\sim)}(t)}{dt} = \delta L_{k-1}^{(L,\sim)}(t) + \alpha f_L^{(L,\sim)} L_k^{(L,\mathrm{V})}(t) - \delta L_k^{(L,\sim)}(t) \quad \text{for } 2 \le k \le n_L, \tag{21v}$$

$$\frac{dL_1^{(L,*)}(t)}{dt} = \alpha\Big(1 - f_L^{(L,\sim)}\Big)L_1^{(L,\mathrm{V})}(t) - \delta L_1^{(L,*)}(t), \tag{21w}$$

and

$$\frac{dL_k^{(L,*)}(t)}{dt} = \delta L_{k-1}^{(L,*)}(t) + \alpha\Big(1 - f_L^{(L,\sim)}\Big)L_k^{(L,\mathrm{V})}(t) - \delta L_k^{(L,*)}(t) \text{ for } 2 \le k \le n_L. \tag{21x}$$

**Dynamics for the number of dead individuals**

After the late infectious phase individuals recover or die. Only symptomatic infections can be lethal. A proportion $f_{\mathrm{Dead}}$ of symptomatic infections is lethal for individuals that were not vaccinated, are unvaccinable or failed to immunize and for which the outcome of the vaccine is still pending. The proportion of lethal individuals is higher among individuals that developed ADE ($f_{\mathrm{Dead}}^{(\mathrm{ADE})} \ge f_{\mathrm{Dead}}$), and lower for individuals with partial immunity ($f_{\mathrm{Dead}}^{(\mathrm{PI})} \le f_{\mathrm{Dead}}$). Individuals that were vaccinated during the fully or late infectious phase with a deleterious outcome also have different proportions of lethal infections, $f_{\mathrm{Dead}}^{(I,*)}$ and $f_{\mathrm{Dead}}^{(L,*)}$, respectively. Hence, the number of dead individuals changes according to

$$\begin{aligned}\frac{dD}{dt} = \delta\Bigg( & f_{\mathrm{Dead}}\Big(f_{\mathrm{Sick}}^{(\mathrm{U},+)} L_{n_L}^{(\mathrm{U},\,+)}(t) + f_{\mathrm{Sick}} L_{n_L}^{(\mathrm{NI})}(t) + f_{\mathrm{Sick}} L_{n_L}^{(\mathrm{V})}(t)\Big) \\ & + f_{\mathrm{Sick}}^{(\mathrm{ADE})} f_{\mathrm{Dead}}^{(\mathrm{ADE})} L_{n_L}^{(\mathrm{ADE})}(t) + f_{\mathrm{Sick}}^{(\mathrm{PI})} f_{\mathrm{Dead}}^{(\mathrm{PI})} L_{n_L}^{(\mathrm{PI})}(t) \\ & + f_{\mathrm{Sick}}^{(I,*)} f_{\mathrm{Dead}}^{(I,*)} L_{n_L}^{(I,*)}(t) + f_{\mathrm{Sick}}^{(L,*)} f_{\mathrm{Dead}}^{(L,*)} L_{n_L}^{(L,*)}\Bigg).\end{aligned} \tag{22}$$

**Dynamics of recovered individuals**

Individuals become permanently immune either by the outcome of the vaccination or by recovering from an infection. All individuals that do not die, recover at the end of the late infectious period and become immune. The number of recovered individuals hence changes according to

$$
\begin{aligned}
\frac{dR}{dt} = &\alpha\Big(f_S^{(R)} S^{(\mathrm{V})}(t) + f_E^{(R)} E_{\mathrm{Sum}}^{(V)}(t) + f_P^{(R)} P_{\mathrm{Sum}}^{(V)}(t) + f_I^{(R)} I_{\mathrm{Sum}}^{(V)}(t) + f_L^{(R)} L_{\mathrm{Sum}}^{(V)}(t)\Big) \\
&+\delta\Bigg(\Big(1 - f_{\mathrm{Sick}}^{(\mathrm{U},+)} f_{\mathrm{Dead}}\Big) L_{n_L}^{(\mathrm{U},\,+)}(t) + L_{n_L}^{(\mathrm{U},\,-)}(t) + \big(1 - f_{\mathrm{Sick}} f_{\mathrm{Dead}}\big) L_{n_L}^{(\mathrm{NI})}(t) \\
&\quad + \big(1 - f_{\mathrm{Sick}} f_{\mathrm{Dead}}\big) L_{n_L}^{(\mathrm{V})}(t) + \Big(1 - f_{\mathrm{Sick}}^{(\mathrm{ADE})} f_{\mathrm{Dead}}^{(\mathrm{ADE})}\Big) L_{n_L}^{(\mathrm{ADE})}(t) \\
&\quad + \Big(1 - f_{\mathrm{Sick}}^{(\mathrm{PI})} f_{\mathrm{Dead}}^{(\mathrm{PI})}\Big) L_{n_L}^{(\mathrm{PI})}(t) + L_{n_L}^{(I,\mathrm{V})}(t) + L_{n_L}^{(I,\sim)}(t) \\
&\quad + \Big(1 - f_{\mathrm{Sick}}^{(I,*)} f_{\mathrm{Dead}}^{(I,*)}\Big) L_{n_L}^{(I,*)}(t) + L_{n_L}^{(I,\mathrm{V})}(t) + L_{n_L}^{(L,\sim)}(t) \\
&\quad + \Big(1 - f_{\mathrm{Sick}}^{(L,*)} f_{\mathrm{Dead}}^{(L,*)}\Big) L_{n_L}^{(L,*)}(t)\Bigg).
\end{aligned} \tag{23}
$$

#### Onset of vaccination campaigns

Vaccination campaigns will be implemented about a year after the outbreak of the pandemic. This requires adaptations in the model. Namely, the vaccination rate must be modelled time dependently. In particular, $\nu$ needs to be replaced by $\nu(t)$ given by

$$
\nu(t) = \begin{cases} 0 \text{ for } t \leq t_{\mathrm{Start}} \\ \nu \text{ for } t > t_{\mathrm{Start}}. \end{cases} \tag{24}
$$

Furthermore the fraction of asymptomatic individuals that get tested positive prior to vaccination, $f_I^{(\mathrm{U},\,+)}$ needs to be replaced by $f_I^{(\mathrm{U},\,+)}(t)$ given by

$$
f_I^{(\mathrm{U},\,+)}(t) = \begin{cases} 0 \text{ for } t \leq t_{\mathrm{Start}} \\ f_I^{(\mathrm{U},\,+)} \text{ for } t > t_{\mathrm{Start}}. \end{cases} \tag{25}
$$

#### Incidence

The incidence $d$-day incidence at time $t$ is defined as the number of new cases that occurred between time $t-d$ and $t$. It is derived as

$$
i_d(t) = \int_{t-d}^{t} \lambda(t) \frac{S(t)}{N} dt. \tag{26}
$$

In the results section of the main manuscript 7-day incidence will be reported, which is obtained by numerical integration.
